# Forebrain Shh overexpression improves cognitive function in a Down syndrome mouse model and euploid littermates

**DOI:** 10.1101/2021.01.18.427185

**Authors:** Feng J. Gao, Donna Klinedinst, Fabian-Xosé Fernandez, Bei Cheng, Alena Savonenko, Benjamin Devenney, Yicong Li, Dan Wu, Martin G. Pomper, Roger H. Reeves

## Abstract

People with Down syndrome (DS) have intellectual disability, early-onset dementia, and cerebellar hypoplasia. Trisomic cerebellar granule cell precursors from Ts65Dn, a mouse model of DS, had a deficit in mitogenic response to Sonic hedgehog (Shh) *in vitro*, and newborn Ts65Dn mice received a single subcutaneous injection of the Shh signaling agonist SAG had normalized cerebellar morphology and improved spatial learning and hippocampal synaptic plasticity at adult. However, cognitive effects of Shh overexpression *in vivo* and where SAG acts to improve cognitive outcomes of trisomy are unknown. Here, we created an inducible human Shh (hShh) knock-in mouse, TRE-bi-hShh-Zsgreen1 (TRE-hShh), expressing dually-lipidated Shh-Np in the presence of transactivator (tTA). Double transgenic mice, Camk2a-tTA;TRE-hShh (Camk2a-hShh) and Pcp2-tTA;TRE-hShh (Pcp2-hShh), increased Shh signaling in forebrain and cerebellum, respectively. Forebrain Shh overexpression normalized hyperactivity, and spatial learning and memory deficits in 3-month-old Ts65Dn, while Shh overexpression in cerebellum had no effect. Further, Camk2a-hShh delayed early-onset severe cognitive impairment in 7-month-old Ts65Dn and enhanced spatial cognition in euploid (Eu) and showed no effect on the longevity of Eu or Ts65Dn, and MRI demonstrated that Pcp2-hShh mitigated disproportionately small cerebellum in Ts65Dn. Finally, Ts65Dn at postnatal day 6 had reduced Gli1 levels in hippocampus and cerebellum, which could be at least partially rescued by Camk2a-hShh and Pcp2-hShh, respectively. Our findings suggest restoration of impaired Shh signaling in forebrain from the perinatal and early postnatal period improves cognitive function.

## Introduction

During gastrulation, both the prechordal plate and the notochord secrete Sonic hedgehog (Shh), which forms a gradient to regulate the central nervous system morphology (Aoto et al., 2009). From embryonic day (E) 8 to E10.5, Shh from the notochord induces primary neurulation to form the neural tube (Goodrich et al., 1996; Placzek, 1995; Ybot-Gonzalez et al., 2007), and the anterior neural tube develops into forebrain, midbrain, and hindbrain that includes cerebellum (Wang et al., 2019). The peak of the forebrain neurogenesis occurs during E10.5-E17, and Shh regulates not only proliferation and differentiation of progenitors in germinal ventricular zone (VZ) (Yabut and Pleasure, 2018) but also cortical interneuron generation and distribution (DeBoer and Anderson, 2017). In the early postnatal and adult forebrain, Shh signaling is also required to maintain neurogenic niches such as dentate gyrus (DG) and subventricular zone (SVZ) (Machold et al., 2003; Palma et al., 2005; Traiffort et al., 2010). During the cerebellum development that occurs mostly between E17.5 and postnatal day (P)15, Shh secreted from Purkinje cells (PCs) regulates granule cell precursor (GCP) proliferation, a process largely determining the cerebellum size and foliation (Dahmane and Ruiz i Altaba, 1999; Wallace, 1999).

Complete knock-out of Shh (Shh^-/-^) is lethal prenatally with a malformed notochord and floorplate and severe tissue patterning defects including cyclopia, the most severe form of holoprosencephaly (HPE) (Chiang et al., 1996; St-Jacques et al., 1998). Haploinsufficiency for Shh has little visible impact on gross phenotype in the mouse (Harfe et al., 2004), but loss of one allele of Shh is sufficient to cause HPE in the human (Roessler et al., 1996). Shh^R34A/K38A^, a double-point mutation in the N-terminal Cardin-Weintraub motif of Shh (KRRHPKK), inhibits the Shh-proteoglycan interaction, and knock-in mice with both endogenous Shh alleles replaced with Shh^R34A/K38A^ show failed Shh accumulation in mitogenic niches, reduced neural precursor proliferation, and growth defects including smaller body and cerebellum (Chan et al., 2009).

Down syndrome (DS), the clinical result of trisomy for human chromosome 21 (HSA21), occurs in about 1 in every 800 babies (Driscoll and Gross, 2009). People with DS have intellectual disability (ID) and a high risk of developing early-onset dementia (Korenberg et al., 1994). HSA21 (GRCh38.p12) has 17 protein coding genes (PCGs) in its short arm (HSA21p) and 213 PCGs in its long arm (HSA21q). Excluding 49 keratin associated protein genes (KRTAPs) that are not expressed in brain, 104, 19 and 37 HSA21q PCGs have respective orthologs in mouse chromosomes (MMU) 16, 17 and 10. Ts65Dn is the most widely used mouse model of DS, which contains an extra freely segregating chromosome with an extra copy of 92 MMU16 non-KRTAP HSA21 orthologs (Gupta et al., 2016) and recapitulates various structural and functional DS brain phenotypes, such as impaired learning and memory, hypocellular forebrain, disproportionally small cerebellum, and early degeneration of forebrain basal neurons (Baxter et al., 2000; Cooper et al., 2001; Escorihuela et al., 1995; Placzek, 1995).

The first direct evidence showing that trisomic progenitors have a Shh response deficit is that isolated Ts65Dn GCPs have a significantly reduced mitogenic response to the dually-lipidated Shh-Np than euploid (Eu) GCPs (Roper et al., 2006). Ts65Dn mice with a single subcutaneous injection of a Shh signaling agonist “SAG 1.1” on the day of birth is sufficient to normalize cerebellar morphology and to improve learning and memory performance in the Morris Water Maze (MWM) performance in adults (Das et al., 2013). Even though forebrain, including cerebral cortex and hippocampus, is considered to be the center for cognitive function, more and more findings such as polysynaptic circuits between cerebellum and forebrain indicate that cerebellum may also play an important role in cognitive function (Buckner, 2013). Here, we generated Eu and Ts65Dn mice with forebrain- or cerebellum-specific Shh overexpression to directly examine the Shh-dose effects on cognitive function.

## Results

### The generation of TRE-hShh knock-in mouse, a new tetracycline inducible transgenic mouse model for human Shh overexpression

Full length Shh (Shh precursor protein) is autoproteolytically cleaved into Shh-N and Shh-C, and fully processed Shh-N (Shh-Np) is dually lipidated, cholesterolized at its C-terminus and palmitoylated at its N-terminus. The dual lipidation is essential to its biological functions such as potency and long-range transport capacity (Chen et al., 2004; Cooper et al., 2003; Lewis et al., 2001; Porter et al., 1996; Zeng et al., 2001). Mice homozygous for Shh mutations that produce loxP or GFP modified Shh-Np are embryonic lethal (Chamberlain et al., 2008; Li et al., 2006). To track the Shh-overexpressing cells without disrupting Shh-Np function *in vivo*, we used pTRE3G-bi vector, containing a doxycycline (Dox) inducible bidirectional promoter capable of driving simultaneous and independent expression of two transgenes, to create the targeting vector, TRE-bi-hShh-Zsgreen1 (TRE-hShh), encoding a Zsgreen1 reporter and a full length hShh cDNA (**Figure S1A**). To avoid the random integration of transgene into the mouse genome that potentially causes disruption of endogenous genes and unexpected phenotypes, we applied Rosa26 TARGATT mice (Tasic et al., 2011) as transgenic recipients using site-specific ϕC31 integrase-mediated transgenesis. Genes of interest in the TRE-hShh vector were flanked by two attB sites to reduce the rate of the transgene integration with bacterial backbone (BB) that could significantly silence adjacent transgene expression (**Figure S1A**).

TRE-hShh targeting vector function was tested by co-transfecting mouse embryonic fibroblasts (MEFs) with CMV-rtTA, which showed that no hShh transcription in the absence of Dox and a prominent induction in its presence (**Figure 1A**). Most full length-hShh was cleaved into hShh-N and hShh-C based on antibodies directed to either the N-terminus (C9C5 and 5H4) or C-terminus (Ab53281) of Shh, while MEFs expressed little or no endogenous Shh protein (**Figures 1B and S1B**). Immunostaining of the co-transfected MEFs with the C9C5 Shh antibody showed that only cells treated with Dox contained Zsgreen1-positive cells and that Zsgreen1 fluorescent intensity was positively correlated with Shh (r=0.68 and p<0.0001, **Figure 1C**).

**Figure 1.**
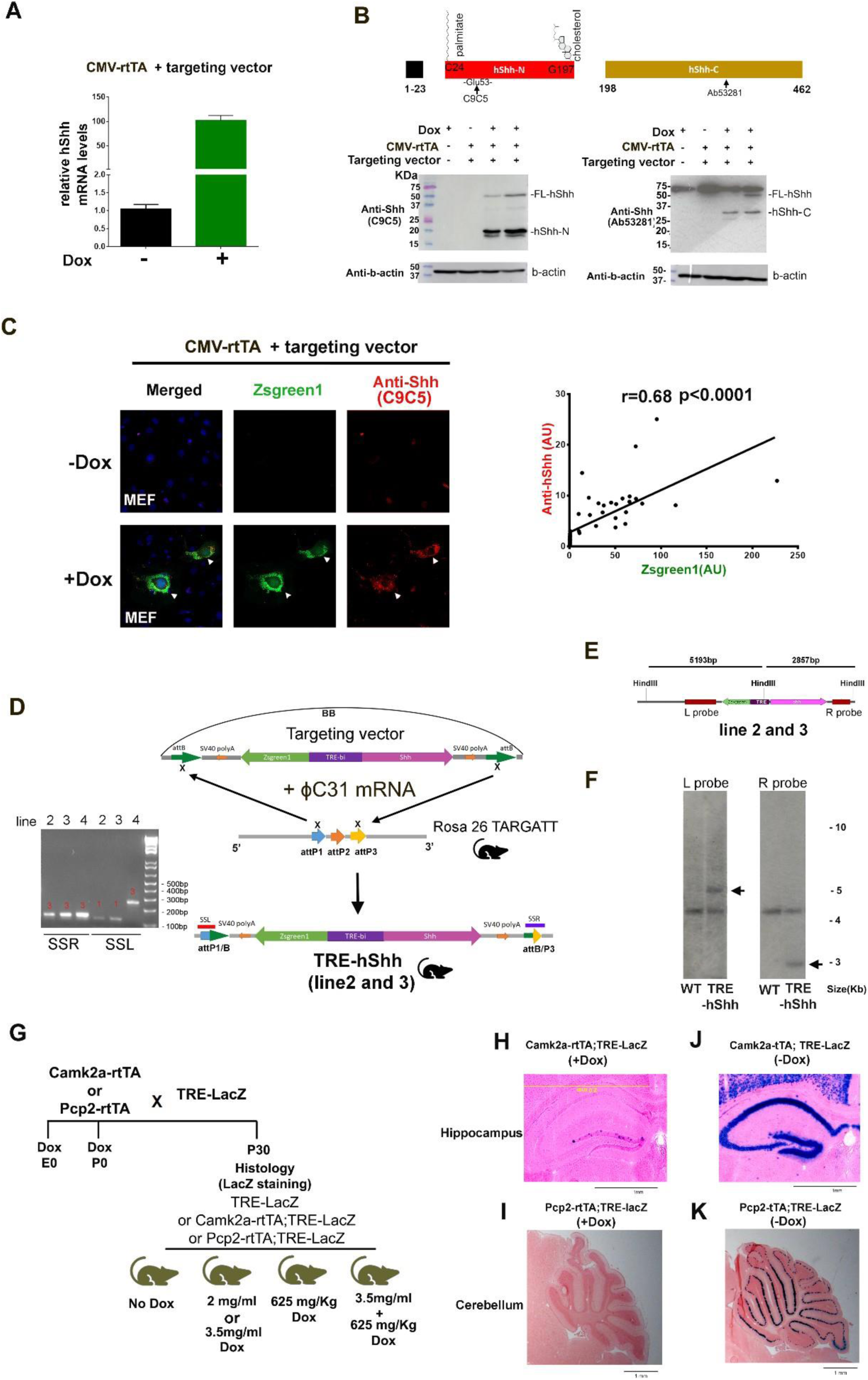
Generation of TRE-hShh transgenic mouse model and verification of rtTA and tTA driver lines for forebrain or cerebellum specific expression. (A) Taqman RT-PCR of MEFs co-transfected with CMV-rtTA and targeting vectors that were treated with or without Dox. Relative levels of hShh mRNA were compared, and beta (b)-actin was used for the RT-PCR normalization. (B) Western blot of MEFs that were co-transfected with CMV-rtTA and targeting vectors, with or without Dox. C9C5 (anti-N-terminus Shh antibody) and Ab53281 (anti-C-terminus Shh antibody) were used, whose targeting sites were shown in the top panel. (C) Immunostaining of MEFs co-transfected with CMV-rtTA and targeting vectors that treated with or without Dox. Representative confocal images of Zsgreen1 and anti-hShh (left panel), and density correlation analysis (right panel). (D) Scheme for generating TRE-hShh transgenic mice through site-specific integration. ϕC31mRNA and the targeting vector were co-injected into the pronucleus of Rosa26 ES with 3 attP sites. PCR of three transgenic lines using site specific primer sets, SSL and SSR. SSL identifies the left junction of attP, and 136bp, 206bp and 282bp PCR products indicate 5’ insertion at attP1, attP2 and attP3, respectively.SSR identifies the right junction of attP, and 225bp and155bp PCR products indicates 3’ insertion at attP2 and attP3, respectively. Transgenic line 2 and 3 had identical attP1/attP3 insertions with gene contents flanked by attB, without the vector’s bacterial backbone (BB). Transgenic line 4 had attP3/attP3 insertion with the whole targeting vector. (E) The scheme to predict southern blot products from TRE-hShh mice (line 2 and 3). HindIII digestion is to produce two fragments. The left fragment is 5193bp that can be detected by L probe, made from PCR products of Tsh3prF2/Tsh3prR3 primers. The right fragment is 2857bp that can be detected by R probe, made from Tsh5prF/Tsh5prR primers. (F) Southern blots of WT and TRE-hShh (line2) using radiolabeled L and R probes. (G) Strategy to obtain TRE-LacZ, Camk2a-rtTA;TRE-LacZ, and Pcp2-rtTA;TRE-LacZ mice for testing the rtTA driver lines. Mice were treated Dox from conception (E0) or from birth (P0) with different Dox concentrations and stained with X-gal at P30, and counter-stained with nuclear fast red, and n≥5 per each group analyzed. (H) Representative images of X-gal stained coronal brain sections from P30 Camk2a-rtTA;TRE-LacZ mice with Dox treatment (625 mg/Kg food pellets plus 3.5 mg/ml in drinking water) from conception. Hippocampus was shown. Images of negative controls including TRE-LacZ(+Dox) and Camk2a-rtTA;TRE-LacZ (-Dox) were shown in Figure S1F. (I) Representative images of X-gal stained sagittal brain sections from P14 mice of Pcp2-rtTA;TRE-LacZ with Dox treatment (625 mg/Kg food pellets plus 3.5 mg/ml in drinking water) from conception. Cerebellum was shown. (J) Representative images of X-gal stained coronal brain sections from P30 Camk2a-tTA;TRE-LacZ mice without Dox treatment. Hippocampus was shown, and more than 3 mice were analyzed. (K) Representative images of X-gal-stained sagittal brain sections from P14 mice of Pcp2-tTA;TRE-LacZ without Dox treatment. Cerebellum was shown, and more than 3 mice were analyzed. See also Figure S1.

The TRE-hShh targeting vector and ϕC31 integrase mRNA were co-injected into the pronucleus of Rosa26 TARGATT mouse ES, which had 3 attP sites within the Rosa26 locus on chromosome 6 to generate TRE-hShh knock-in mouse. PCR for hShh and Zsgreen1 found 3 of 15 pups with the targeting vector insertion. With commercially available site-specific primer sets, SSL and SSR, we identified that founder line 2 and 3 had identical attP1/attP3 transgene insertions without BB (**Figure 1D**), and that line 4 had the whole plasmid (with BB) inserted at attP3 site (**Figure S1C**). Southern blots of wildtype (WT) and the three transgenic lines validated respective transgene insertions (**Figures 1E-F, S1D-E**). All three lines were backcrossed into C57BL/6J (B6) for > 6 generations, and mice heterozygous or homozygous for TRE-hShh were viable and fertile.

### Camk2a-tTA and Pcp2-tTA drive TRE-LacZ expression in forebrain and cerebellum, respectively

The stimulation of the Shh pathway by treatment of Ts65Dn mice with SAG at P0 restores cerebellar morphology and improves cognitive function in adult (Das et al., 2013). Since SAG can reach every tissue and exert its effects in Shh-responsive cells ubiquitously, it is unclear from this result whether improved cognitive function is due to stimulation of the Shh pathway in hippocampus, or cerebellum, or possibly elsewhere in the brain. Thus, we sought to express hShh beginning in the early postnatal period in hippocampus or cerebellum. The Camk2a promoter is restrictively active in the forebrain including hippocampus (Tsien et al., 1996), beginning at P1 and its activity increases significantly by P5 (Bayer et al., 1999). The Pcp2 promoter drives expression in a Purkinje cell (PC) specific manner from E17.5 (Lewis et al., 2004).

We tested both the Tet-on driver lines, Camk2a-rtTA (Mansuy et al., 1998) and Pcp2-rtTA (unpublished), and the Tet-off driver lines, Camk2a-tTA (Mayford et al., 1996) and Pcp2-tTA (Zu et al., 2004) using a TRE-LacZ reporter line (Furth et al., 1994). In the Tet-on system, neither Camk2a-rtTA nor Pcp2-rtTA effectively induced TRE-LacZ expression, even with continuous Dox exposure from conception (**Figures 1G-I, S1F, and Table S1**). In contrast, Camk2a-tTA and Pcp2-tTA drove TRE-LacZ expression in a forebrain- and cerebellum-specific manner, respectively (**Figures 1J-K**).

### Camk2a-tTA;TRE-hShh mice express dually-lipidated hShh-Np

We crossed B6.Camk2a-tTA with all three B6.TRE-hShh lines (**Figure 2A**). Double transgenic mice, Camk2a-tTA;TRE-hShh (Camk2a-hShh), from line 2 and 3 had Zsgreen1 expression pattern similar to LacZ expression of Camk2a-tTA;TRE-LacZ, while Camk2a-hShh from line 4 had little or no expression in forebrain (**Figure S2A**). We used the line 2 for subsequent experiments. Both recombinant hShh-Np purified from HEK293 cells and the lysates of MEF cells that were co-transfected with CMV-rtTA and TRE-hShh plus Dox showed three ∼20 kDa bands in SDS-PAGE visualized by Coomassie blue staining and Western blot, and the highest molecular weight band was the most prominent (**Figures 2B and S2B**). Human Shh-N, the C24-G197 of Shh precursor protein (462 amino acids), has a theoretical molecular weight (MW) of 19.56 kDa prior to post-translational modifications, of which palmitoylation at C24 adds ∼238 Da and cholesterol addition at G197 is ∼369 Da, and contains almost identical sequences to mouse Shh-N, the only amino acid difference is that S44 of human Shh-N becomes T44 in mouse Shh-N (**Figure S2C**). MALDI-TOF mass spectrometry of the recombinant hShh-Np showed the peak at ∼20.173 kDa (**Figure 2C**), which matches the theoretical MW of the dually-lipidated hShh-Np form. Unlike MEF with Shh-transient transfection or recombinant hShh-Np, both Camk2a-hShh and control mice only showed the dually-lipidated Shh-Np form based on Western blot analysis (**Figure 2B)**, indicating a more stringent quality control of Shh processing *in vivo*.

**Figure 2.**
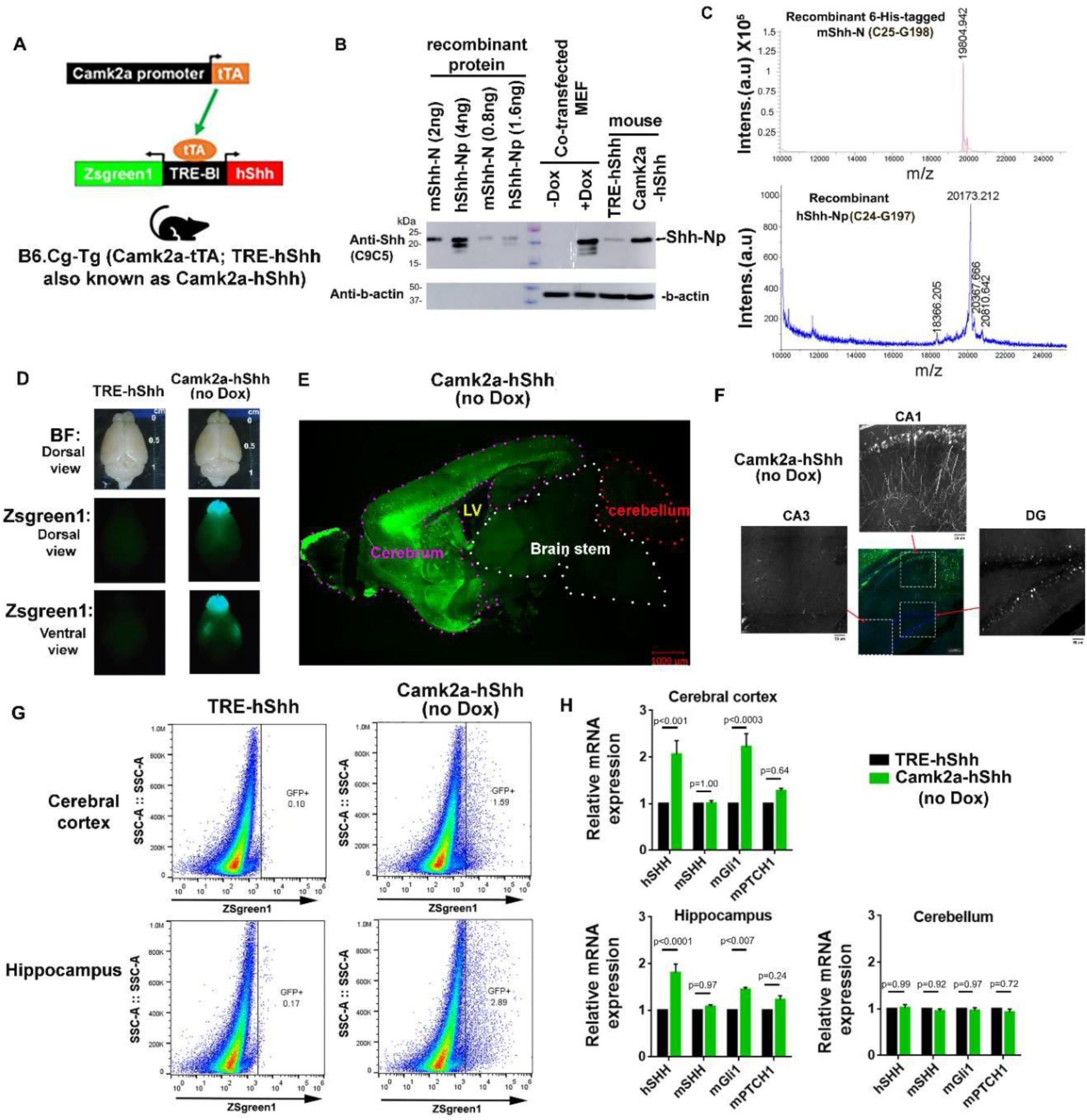
Camk2a-tTA induces dually-lipidated hShh-Np expression from TRE-hShh that enhances Shh signaling in forebrain. (A) Scheme of generating Camk2a-tTA;TRE-hShh mice. When Camk2a promoter is active, tTA binding to TRE-hShh that drives Zsgreen1 and hShh expression simultaneously. (B) Western blot of purified recombinant hShh-Np, 6xHis tagged mShh-N, lysate of MEF co-transfected of CMV-rtTA and the targeting vectors, and mouse cortex lysate of TRE-hShh and Camk2a-hShh that were probed with anti-hShh (C9C5) and anti-b-actin. (C) MALDI-TOF MS analysis of purified recombinant protein, 6xHis tagged mShh-N (top) and hShh-Np (bottom). (D) Two-month-old mouse brains of TRE-hShh and Camk2a-hShh under bright-field (BF) or GFP channel. (E) Representative tile scan confocal images of Zsgreen1-positive cells in a sagittal brain section of 2-month-old Camk2a-hShh. (F) Zsgreen1-positive cells in Camk2a-hShh hippocampus. (G) FACS of dissociated neurons from cerebral cortex or hippocampus of 2-month-old TRE-hShh and Camk2a-hShh mice. The same gate setting was used to separate Zsgreen1-negative and Zsgreen1-positive neurons for all samples, and n=2 per group. (H) Taqman RT-PCR of key Shh signaling transcripts, hShh, mShh, mGli1, and mPTCH1 from cortex, hippocampus, and cerebellum. TRE-hShh and Camk2a-hShh were compared (n=5 per group), and b-actin was used for the RT-PCR normalization. Data were represented as mean ± SEM and analyzed using two-way ANOVA and Sidak’s multiple comparisons test. See also Figure S2 and Video S1 and S2.

### Camk2a-tTA;TRE-hShh has enhanced Shh signaling in forebrain

The forebrain of P1 Camk2a-hShh live pups without Dox treatment was GFP-positive that could be identified by GFP flashlight (Nightsea) (**Figure S2E**). In 2-month-old mice, Camk2a-hShh had Zsgreen1-positive forebrain, and no general brain morphology difference was found between TRE-hShh and Camk2a-hShh (**Figure 2D**). Sagittal brain sections of Camk2a-hShh showed that forebrain structures including olfactory bulb, cortex, hippocampus, and basal ganglia were Zsgreen1-positive, while cerebellum and brain stem were Zsgreen1-negative (**Figure 2E**). In the hippocampus, Zsgreen1-positive neurons were densely packed in dentate gyrus (DG) and CA1 but sporadic in CA3 (**Figure 2F, Video S1 and S2**). Using a conservative gate setting for GFP-positive cells, fluorescence activated cell sorting (FACS) of Camk2a-hShh showed ∼1.6% Zsgreen1-positive cells in the cortex and ∼2.9% in the hippocampus (**Figure 2G**).

Expression of canonical Shh signaling pathway genes *Shh, Patched1 (PTCH1),* and *Gli1* was compared between TRE-hShh and Camk2a-hShh mice at 2-month-old (**Figure 2H**). In cortex or hippocampus, hShh transcripts in Camk2a-hShh levels were twice as high as those in TRE-hShh. Based on FACS quantification of Zsgreen1-positive cells, we deduced that the presence of Camk2a-tTA induced TRE-hShh expression by at least 50 times. Mouse Gli1 (mGli1), a sensitive readout of active Shh signaling, was significantly increased in Camk2a-hShh (p<0.0003 and p<0.007 in cortex and hippocampus, respectively). As endogenous mShh transcripts were not changed (p=1.0 in cortex and p=0.97 in hippocampus), we attributed Shh signaling activation in forebrain of Camk2a-hShh to hShh expression. Together, these results show that Camk2a-hShh enhanced Shh signaling in forebrain through overexpression of the dually-lipidated Shh-Np form as endogenous Shh gene does, which is essential to study Shh dosage effects as the expression of unmodified Shh-N may have dominant negative effects *in vivo* (Huang et al., 2007; Lewis et al., 2001).

### Pcp2-tTA;TRE-hShh shows PC-specific hShh expression and enhances Shh signaling in cerebellum

Sagittal brain sections of P0, P6 and P14 PCP2-tTA;TRE-LacZ mice were analyzed by X-gal staining (**Figure 3A**). Multiple layers of small and low-intensity of LacZ-positive cells were detected at P0, which matches PC aggregation in a primordial cortical layer of cerebellum at the stage (Rahimi-Balaei et al., 2018; Sergaki and Ibanez, 2017). The size and intensity of LacZ-positive cells increased and formed a single layer by P14. B6.Pcp2-tTA mice were crossed with B6.TRE-hShh to generate Pcp2-tTA;TRE-hShh (Pcp2-hShh, **Figure 3B**). Zsgreen1 expression was restricted in the cerebellum of 2-month-old Pcp2-hShh mouse (**Figure 3C**), and Calbindin1 immunostaining confirmed that only PCs were Zsgreen1-positive (**Figure 3D and Video S3**). FACS of Pcp2-hShh showed Zsgreen1-positive cells in cerebellum but not in cortex or hippocampus (**Figures 3E and S3**). Comparing to TRE-hShh, Pcp2-hShh had significantly increased mGli1 transcript levels in cerebellum, with the higher induction at P6 than P60 (**Figure 3F)**, which was likely due to more Shh responding cells such as GCPs in the developing postnatal cerebellum than the developed adult cerebellum. Camk2a-tTA and Pcp2-tTA were able to drive TRE-LacZ expression in P6 forebrain and cerebellum, respectively (**Figure 3G**), supporting that Camk2a-tTA and Pcp2-tTA are capable of inducing TRE promoter in forebrain or cerebellum from the perinatal/early postnatal period.

**Figure 3.**
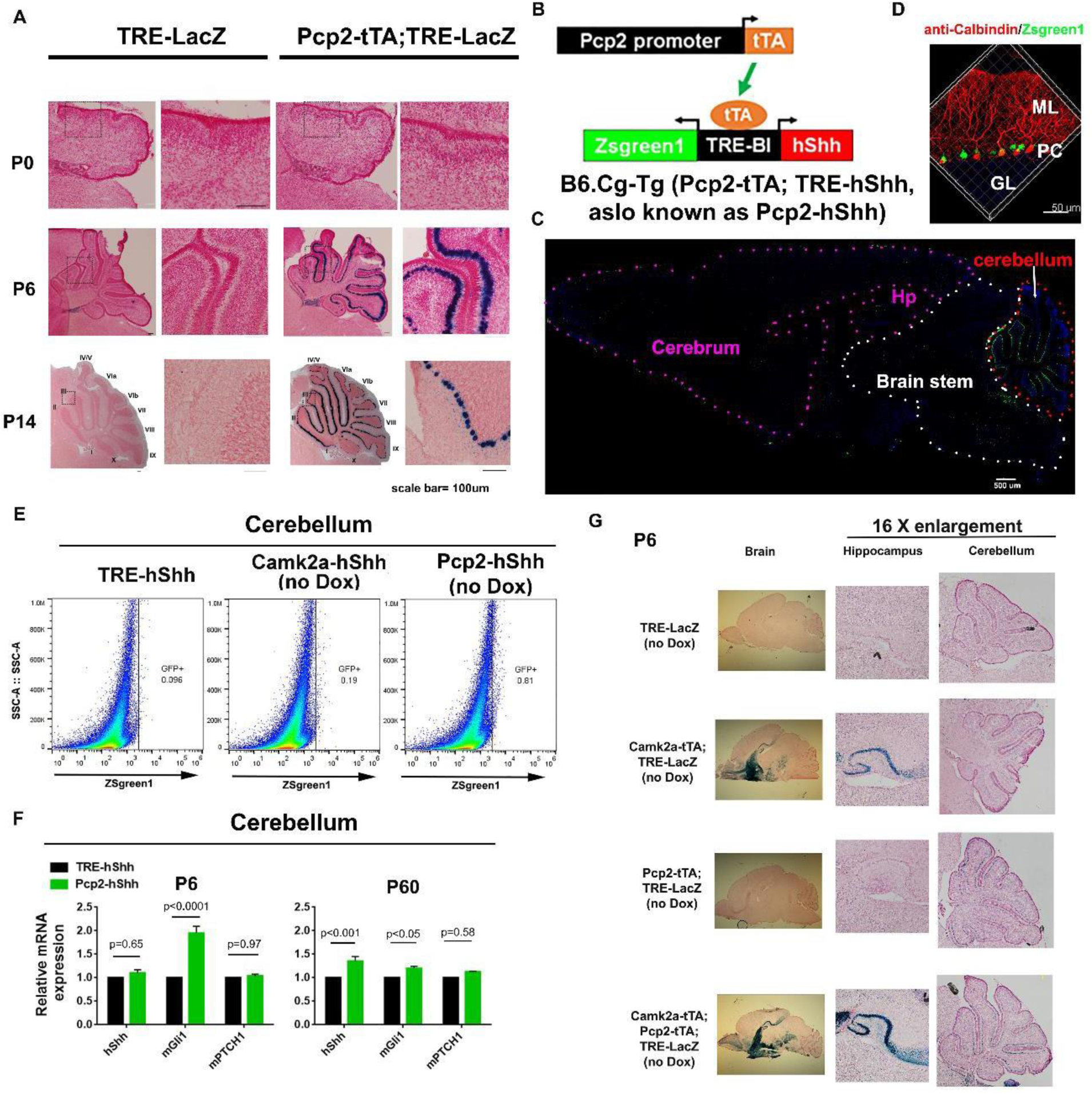
Pcp2-hShh induces PC-specific hShh overexpression and enhances Shh signaling in cerebellum. (A) X-gal staining of sagittal brain sections of TRE-LacZ and PCP2-tTA;TRE-LacZ at P0, P6, and P14, and counter-stained with nuclear fast red. Cerebellum areas were compared. (B) Scheme of generating the B6.Pcp2-tTA;TRE-hShh double transgenic mouse. (C) Representative tile scan confocal images of Zsgreen1-positive cells in 2-month-old Pcp2-hShh mouse brain. (D) Immunostaining of Pcp2-hShh sagittal brain sections for Calbindin (red). Granular layer (GL), PCs, and molecular layer (ML) of cerebellar cortex were shown. (E) FACS of cerebellums from TRE-hShh, Camk2a-hShh, and Pcp2-hShh mice. (F) Taqman RT-PCR for hShh, mGli1 and mPTCH1 from TRE-hShh and PCP2-hShh cerebellum at P6 and P60 (n=5 per group). Data were represented as mean ± SEM and analyzed by repeated two-way ANOVA and Sidak’s multiple comparisons test. (G) Brain sagittal sections of P6 Littermates of TRE-LacZ, Camk2a-tTA;TRE-LacZ, Pcp2-tTA;TRE-LacZ, Camk2a-tTA;Pcp2-tTA;TRE-LacZ, which were offspring of Pcp2-tTA and Camk2a-tTA;TRE-LacZ, were analyzed by X-gal staining. The sections were counter-stained with nuclear fast red. See also Figure S3 and Video S3.

### Camk2-hShh but not Pcp2-hShh normalizes hyperactivity in Ts65Dn

All single-transgenic lines, Camk2a-tTA, Pcp2-tTA, and TRE-hShh, were inherited according to the Mendelian ratio (**Figure S4A**). When breeding for double-transgenic lines, populations of both Camk2a-hShh and Pcp2-hShh were greater than their TRE-hShh littermates (p=0.05 and p<0.03, respectively, **Figure S4B**). Camk2a-hShh mice with Dox from conception to E17 had significantly lower Zsgreen1 expression than ones without Dox treatment at P40 (**Figure S4C**), consistent with previous demonstrations that Tet-off system treated with Dox during embryonic development prevents transgene from full expression in adult (Krestel et al., 2001).

In this study, both “forebrain-cohort” and “cerebellum-cohort” were generated with no Dox treatment to use the Tet-off system to its full potential. Behavioral tests including open field, visual discrimination, and Morris water maze (MWM) started at 3-month-old (**Figure 4A**). The forebrain cohort consisted of 14-16 males in each of four groups, Eu;TRE-hShh (Eu control), Eu;Camk2a-hShh (Eu with forebrain hShh overexpression), Ts65Dn;TRE-hShh (Ts65Dn control), and Ts65Dn;Camk2a-hShh (Ts65Dn with forebrain hShh overexpression). The cerebellum cohort consisted of 18-20 males in each of four genotypes, Eu;TRE-hShh (Eu control), Eu;Pcp2-hShh (Eu with cerebellum hShh overexpression), Ts65Dn;TRE-hShh (Ts65Dn control), and Ts65Dn;Pcp2-hShh (Ts65Dn with cerebellum hShh overexpression). In 3-month-old mice, Camk2-hShh expressed significantly more Shh-Np in forebrain than TRE-hShh controls **(Figure S4D)**, and Pcp2-hShh cerebellum had significantly more Gli1 protein than controls (**Figure S4E**). Shh overexpression in forebrain or cerebellum had no significant effect on mouse body weight (**Figure 4B**).

**Figure 4.**
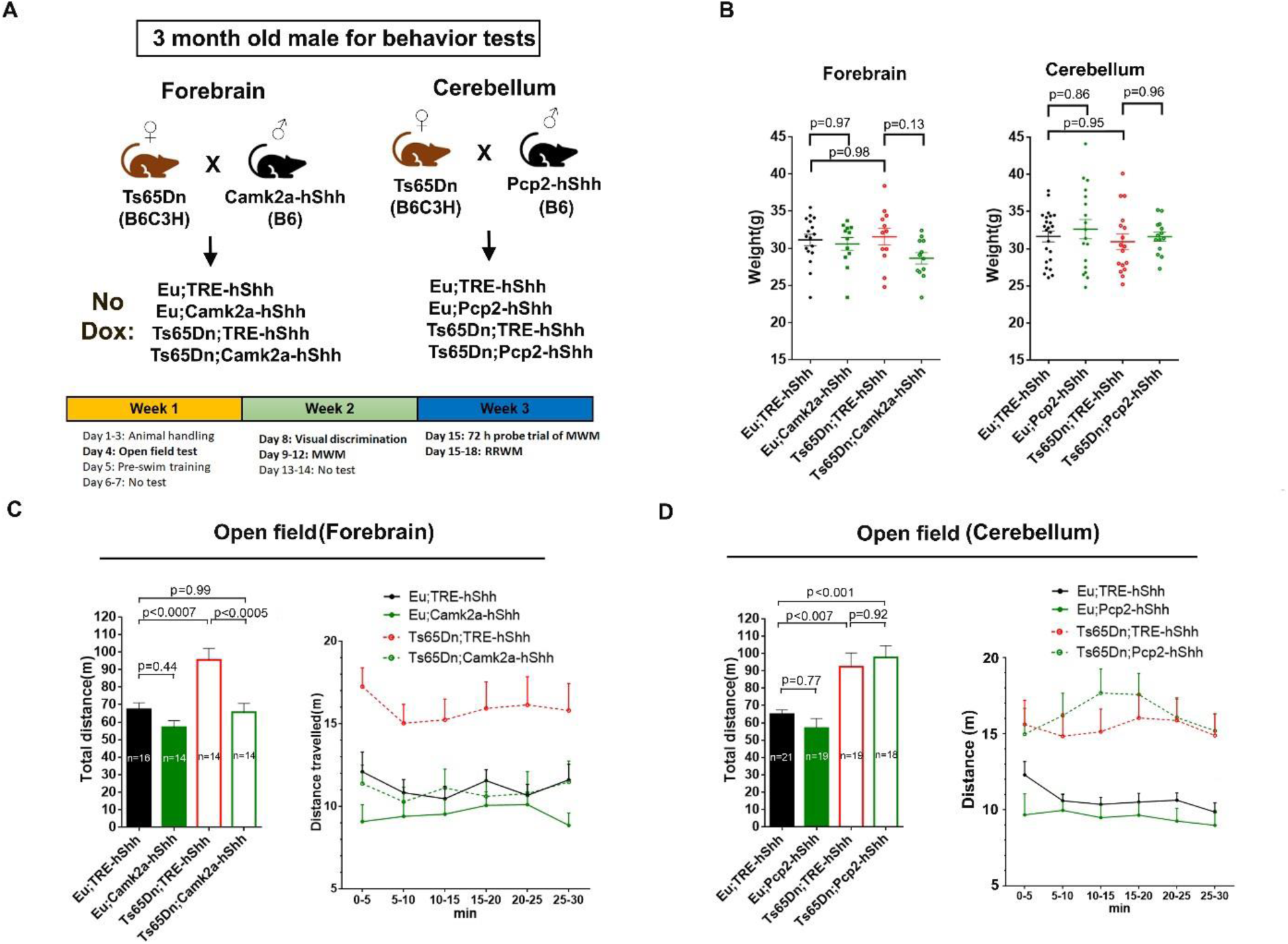
Camk2-hShh but not Pcp2-hShh completely normalizes hyperactivity in Ts65Dn. (A) Scheme for generating the forebrain cohort (n=14-16 per group) and the cerebellum cohort (n=18-21 per group) for behavioral tests. All animals were not treated with Dox. The timeline of each behavioral test was shown. (B) Body weight of different transgenic mice at 3-month-old. (C) 30 min open field tests of the forebrain cohort. Total distance travelled in 30 min (left) and distance travelled in each of 5-min-bin (right) were shown (n=14-16 per group). (D) 30 min open field tests of the cerebellum cohort. Total distance travelled in 30 min (left) and distance travelled in each of 5-min-bin (right) were shown (n=18-21 per group). Data were represented as mean ± SEM. Data were analyzed by one-way ANOVA and Tukey’s multiple comparisons test (B). Total distance was travelled during 30 min (the left panel of C and D), which were analyzed by one-way ANOVA and Tukey’s multiple comparisons test. Distance was travelled in each of 5-min-bin (the right panel of C and D), which were analyzed by two-way RM ANOVA and Tukey’s multiple comparisons test. See also Figure S4.

Hyperactivity in Ts65Dn has been observed repeatedly since the model was established (Escorihuela et al., 1995; Faizi et al., 2011; Fortress et al., 2015). In a 30 min novel open field paradigm, forebrain Shh overexpression in Ts65Dn completely normalized its hyperactivity based on the total distance travelled (p<0.0003 for Eu;TRE-hShh VS Ts65Dn;TRE-hShh, and p=0.95 for Eu;TRE-hShh VS Ts65Dn;Camk2a-hShh, **Figure 4C**). In the cerebellum cohort, Shh overexpression did not normalize hyperactivity of Ts65Dn (p<0.005 for Eu;TRE-hShh VS Ts65Dn;TRE-hShh, and p<0.0007 for Eu;TRE-hShh VS Ts65Dn;Pcp2-hShh **Figure 4D**).

### Camk2-hShh but not Pcp2-hShh improves learning and memory in Eu and Ts65Dn

To test whether Shh overexpression has any effect on the visual ability and goal-directed behaviors, mice were tested in six-trial visual discrimination (VD), a non-spatial learning water maze with a cued hidden platform (**Figure S5A**). Ts65Dn performed slightly worse than Eu (p=0.08 in forebrain-cohort and p=0.06 in cerebellum-cohort), and hShh overexpression in forebrain or cerebellum had no significant effect on the VD performance (**Figure S5B**).

To assess spatial learning and memory, all mice were tested in MWM (**Figure S6A**). As Ts65Dn swam faster than Eu in acquisition trials (p<0.0001, **Figure S6B**), the performance of acquisition trials was reported as escape distance rather than latency. All groups reduced escape distance to find the hidden target platform as training progressed, but the improvement was significantly slower in Ts65Dn than Eu (p<0.0001, **Figure 5A**). In the forebrain-cohort, hShh overexpression significantly reduced escape distance in Ts65Dn (p<0.02). In the cerebellum-cohort, hShh overexpression had no significant effect on the escape distance of Ts65Dn or Eu (p=0.1 and p=0.65, respectively)

**Figure 5.**
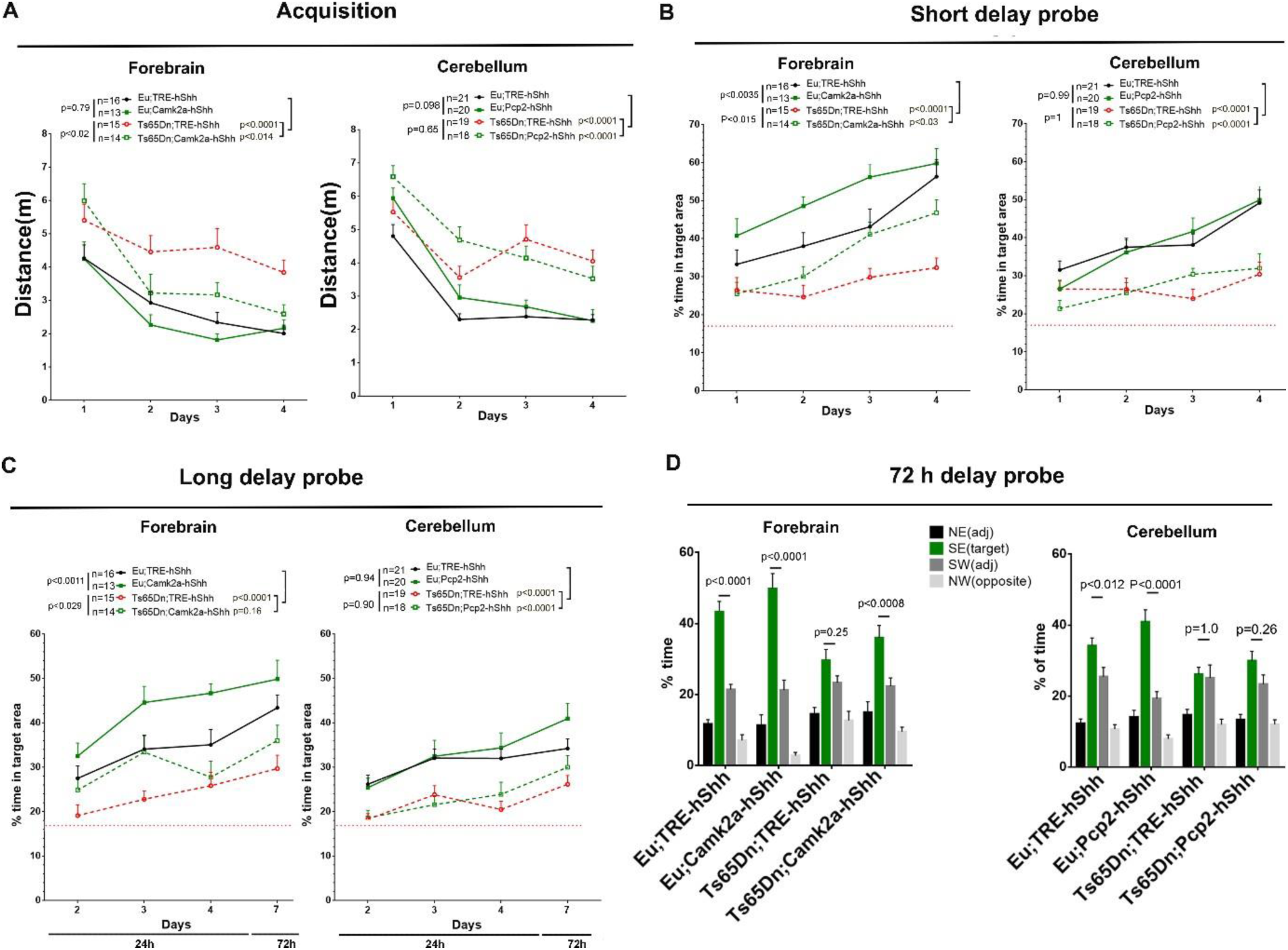
Camk2-hShh but not Pcp2-hShh improves learning and memory in MWM. (A) Escape distance in acquisition trials from training day 1 to 4. (B) Short (30 min) delay probe trials from training day (TD) 1 to 4. (C) Long delay probe trials. TD2-TD4 were 24 h delay probe trials, and TD7 were 72 h delay probe trials. (D) The quadrant preference analysis of the 72 h long delay probe trials. Data were represented as mean ± SEM and analyzed by two-way ANOVA and Tukey’s multiple comparisons tests. N=13-16 per group in the forebrain cohort, and n=18-21 per group in the cerebellum cohort. See also Figure S5 and S6.

Four-day overall performance in short delay probe trials (**Figure 5B)** showed that Ts65Dn lagged significantly behind Eu (p<0.0001), and that forebrain hShh overexpression significantly improved the performance in both Eu and Ts65Dn (p<0.0035 and p<0.015, respectively) and it also mitigated the deficiency caused by trisomy (p<0.0001 for Eu;TRE-hShh VS Ts65Dn;TRE-hShh, p<0.03 for Eu;TRE-hShh VS Ts65Dn;Camk2a-hShh). In the cerebellum-cohort, hShh overexpression had no significant effect on Eu or Ts65Dn (p=0.99 and p=1.0, respectively).

In four-day overall performance in long delay probe trials (**Figure 5C**), forebrain hShh overexpression significantly enhanced the performance in both Eu and Ts65Dn (p<0.0011 and p<0.029, respectively) and rescued the trisomy caused deficiency (p<0.0001 for Eu;TRE-hShh VS Ts65Dn;TRE-hShh, p=0.16 for Eu;TRE-hShh VS Ts65Dn;Camk2a-hShh). The quadrant preference analysis of 72 h long delay probe trials showed that forebrain Shh overexpression significantly improved the performance in Ts65Dn (**Figure 5D**). Cerebellum hShh overexpression had no significant effect on the performance of long delay probe trials of Eu or Ts65Dn (p=0.94 and p=0.90, respectively).

### Camk2a-hShh improved the ability to forget non-essential memory

To assess the effects of forebrain hShh overexpression on the integrity of learning and memory (i.e. inhibition of non-essential memory and retention of essential spatial reference memory), the forebrain cohort was tested in repeated reversal water maze (RRWM, **Figure S7A**) immediately following the 72 h probe trial of MWM. In reversal session 1 (reversal (R) day 1 and R day 2), the hidden platform was moved from SE to NW, and the platform was then relocated to SW in reversal session 2 (R day 3 and R day 4). Thus, the old platform position “SE” is the non-essential memory in RRWM, and the spatial reference is the essential memory as RRWM has the same spatial reference as MWM.

Firstly, to quantify the ability to forget the non-essential memory, we compared % of time spent in SE target area in long delay probe before RRWM training (trial 1 of R day1), with that of after 1 day training (trial 1 of R day2), and with that after 2 days training (trial 1 of R day3, **Figure 6A**). After 1 day RRWM training, the average SE memory reduction were 40% in Eu;TRE-hShh (p=0.0002), 61% in Eu;Camk2a-hShh (p<0.0001), 4% in Ts65Dn;TRE-hShh (P=0.93), and 23% in Ts65Dn;Camk2a-hShh (p=0.077). Secondly, probe trials showed that forebrain Shh expression significantly increased time spent in new target areas (p=0.024 in Eu, and p=0.035 inTs65Dn, **Figure 6B**), indicating a better retention of the essential memory, the spatial reference.

**Figure 6.**
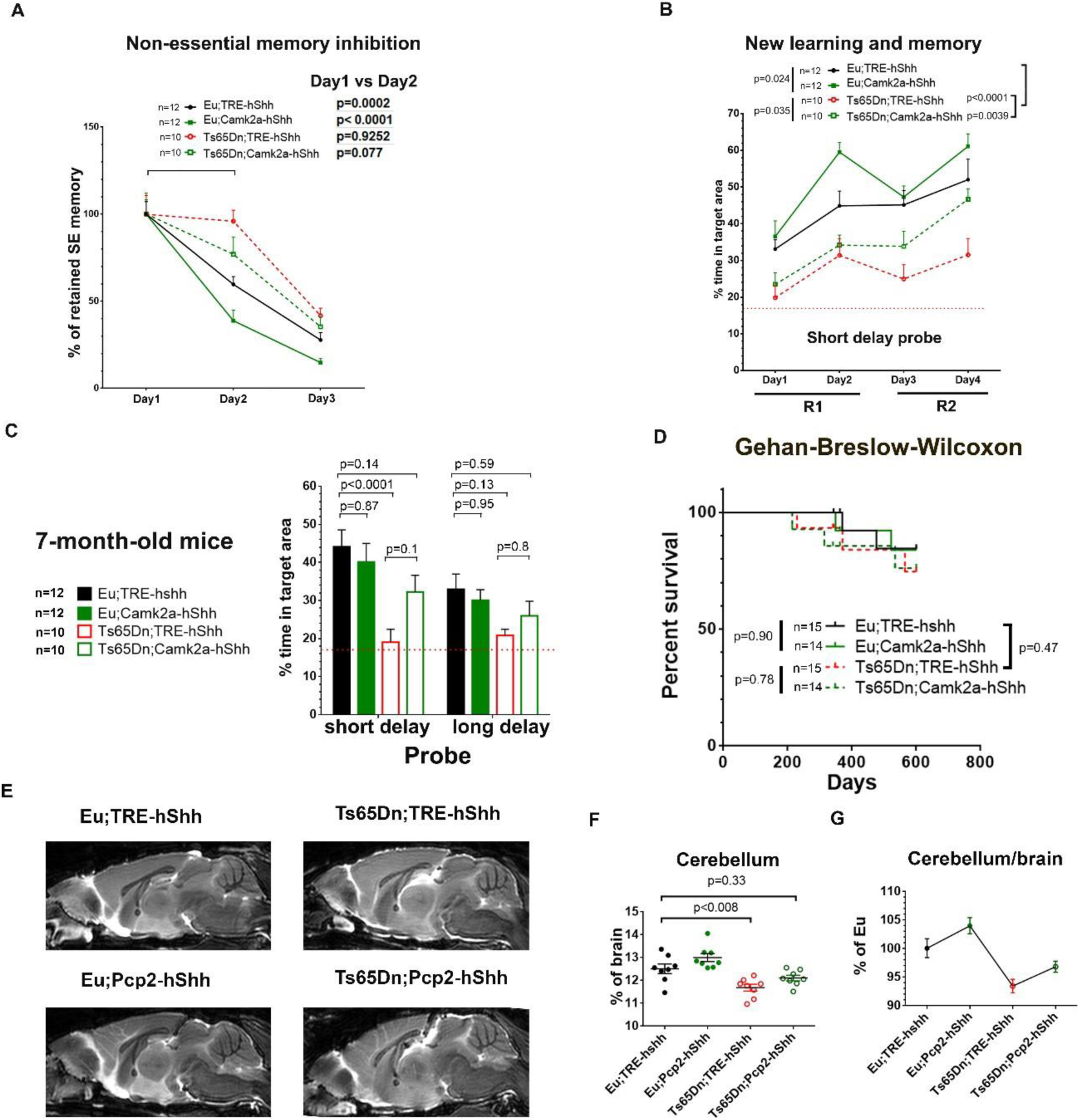
RRWM and survival analysis of forebrain-cohort, and MRI of cerebellum-cohort. (A) RRWM of the forebrain cohort. Long delay probe trials for the old platform position “SE” from day1 to day3, and n=10-12 per group. (B) RRWM short delay probe trials for new platform positions. R, reversal session. (C) The forebrain cohort were re-tested in MWM at 7-month-old (n=10-12 per group). (D) Survival analysis of the forebrain cohort aged until 600 days old (n=14-15 per group). (E) Representative high-resolution 3D T2-weighted MRI of the 14-month-old cerebellum cohort. The mid-sagittal brain images were shown. (F) Statistical analysis of the cerebellum/brain volume ratio of the 14-month-old cerebellum cohort based on the MRI (n = 8 per group). (G) The cerebellum/brain ratio of other groups relative to Eu controls based on the MRI. Data were represented as mean ± SEM. Data were analyzed by two-way ANOVA and Tukey’s multiple comparisons test (A-C), and by Gehan-Breslow-Wilcoxon test (D), and by two-way ANOVA and Tukey’s multiple comparisons test (F-G). See also Figure S7 and S8.

### Camk2a-hShh delays early-onset severe cognitive impairment in 7-month-old Ts65Dn

People with DS have a high risk of developing early onset dementia, and Ts65Dn mice fail to acquire any spatial learning and memory in MWM at ∼ 6-month-old (Brose et al., 2018; Brose et al., 2019). To test whether the cognitive improvement by forebrain hShh overexpression is long-lasting, the forebrain cohort were re-tested in MWM with different spatial references at ∼7-month-old. The short (30 min) delay probe trials of day 4 training showed that the average percentage of time spent in target area was 19% in Ts65Dn;TRE-hShh (close to the 17% chance level), 32% in Ts65Dn;Camk2a-hShh, and 41% in Eu;TRE-hShh, and forebrain hShh overexpression mitigated the trisomy caused deficiency (p<0.0001 for Eu;TRE-hShh VS Ts65Dn;TRE-hShh, and p=0.14 for Eu; TRE-hShh VS Ts65Dn;Camk2a-hShh, **Figure 6C**).

To evaluate side effects of chronic forebrain hShh overexpression, the forebrain cohort (14-15 per group) was aged until 600 days old for survival analysis. The Gehan-Breslow-Wilcoxon analysis, giving more weight to early deaths, showed that the mortality rate of Ts65Dn was slightly higher than Eu (p=0.47), and that chronic hShh forebrain overexpression showed no significant effect on longevity (p=0.90 in Eu, and p=0.78 in Ts65Dn, **Figure 6D**).

### Pcp2-hShh mitigates disproportionately small cerebellar volumes in Ts65Dn

Individuals with DS have a disproportionately small cerebellum (Aylward et al., 1997), which is present in various DS mouse models such as Ts65Dn, Ts1Cje, and TcMAC21 (Kazuki et al., 2020; Olson et al., 2004). Postnatal cerebellar development is critical (Fujishima et al., 2012; Martinez et al., 2013), which is driven in part by Shh expression from PCs. The proliferation of both Ts65Dn and Eu cerebellar GCPs show a linear dose response to Shh, but significantly lower response to the same Shh concentration in trisomy (Roper et al., 2006). SAG treatment of P0 Ts65Dn normalizes cerebellar morphology in adult (Das et al., 2013).

To quantify the cerebellar volume changes, high-resolution 3D T2-weighted MRI was performed on the 14-month-old cerebellum-cohort (**Figure 6E**). Two-way ANOVA analysis showed that the trisomic effect significantly reduced the cerebellum/brain ratio (p<0.0001), and that cerebellum hShh overexpression significantly increased the cerebellum/brain ratio (p=0.01). Tukey’s post-hoc multiple comparisons showed that Pcp2-hShh mitigated the trisomy-caused disproportionately small cerebellum (p<0.008 for Eu;TRE-hShh VS Ts65Dn;TRE-hShh, and p=0.33 for Eu; TRE-hShh VS Ts65Dn;Pcp2-hShh, **Figures 6F-G**). Relative to the average cerebellum/brain ratio of Eu;TRE-hShh (100%), it was 104.0% in Eu;Pcp2-hShh, 93.5% in Ts65Dn;TRE-hShh, and 96.9% in Ts65Dn;Pcp2-hShh.

### Ts65Dn has reduced Gli1 activities in both hippocampus and cerebellum at P6

Shh signaling is essential to proliferation and differentiation of neuronal progenitors (NPs) during the development of forebrain (DeBoer and Anderson, 2017; Yabut and Pleasure, 2018) and cerebellum (Dahmane and Ruiz i Altaba, 1999; Wallace, 1999), and it is also required to maintain adult neural stem cells in neurogenic niches (Machold et al., 2003; Palma et al., 2005; Traiffort et al., 2010). Shh signal transduction in Shh-responding cells including NPs is through activation of one or more of three Gli transcription factors that could regulate expression of G1 cell cycle proteins such as cyclin D and N-Myc and the antiapoptotic protein Bcl-2, which at least partially is responsible for controlling NP fate (Cayuso et al., 2006; Yao et al., 2016). To track changes in Shh-responding cells in brain, we used a Gli1-LacZ reporter mouse, which has LacZ inserted into the first coding exon (exon 2) of Gli1 (Bai et al., 2002). Thus, LacZ expression is mimicking the endogenous Gli1 activity.

In hippocampus, LacZ-positive cells were mainly located in the polymorphic layer and sub-granular layer of DG, and the population of LacZ-positive cells were significantly reduced from P6 to P21 to P90 (**Figure 7A**). To define Gli1-positive cells in cerebellum, brain sections of P6, P21 and P90 Gli1-LacZ mice were costained by X-gal and anti-Calbindin immunostaining **(Figure 7B-C)**. At P6, most LacZ-positive cells were in outer external granular layer (oEGL) that contained GCPs, and a small population of LacZ-positive cells were located beneath Purkinje cell layer (bPCL). At P21, LacZ-positive oEGL was completely disappeared, while bPCL was still LacZ-positive and merging into PCL. At P90, bPCL was completely merged with PCL and contained LacZ-positive and calbindin-negative cells. Previous studies linked Shh responding cells in adult PCL to Bergmann glia, a multi-functional astrocyte retaining neural precursor properties (Koirala and Corfas, 2010; Traiffort et al., 1999).

**Figure 7.**
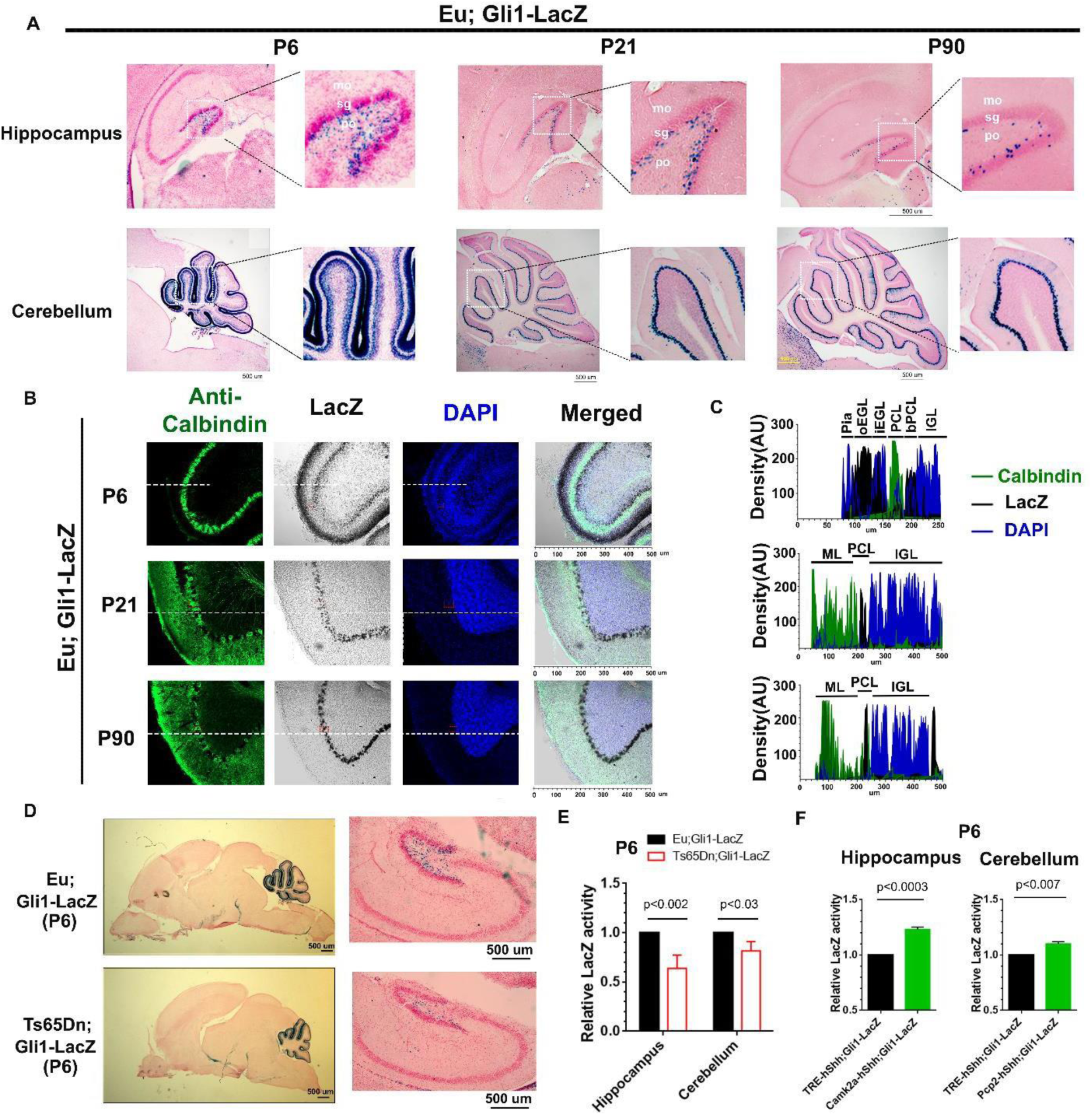
Ts65Dn has reduced Gli1 activities in both hippocampus and cerebellum at P6. (A) X-gal staining of sagittal brain sections of Eu; Gli1-LacZ at P6, P21 and P90. Hippocampus and cerebellum areas were shown. mo, molecular layer; sg, subgranular zone; po, polymorphic layer. Scale bar=500 um. (B) Representative images of P6, P21 and P90 of Eu; Gli1-LacZ cerebellum costained for LacZ (black) and Calbindin (green), and nuclear DNA was stained with DAPI (blue). (C) Colocalization analysis of white dash labeled areas in (B). Six distinguished layers found at P6, from outermost to innermost: Pia, high DAPI, LacZ and Calbindin negative; oEGL, low DAPI, high LacZ positive, and Calbindin negative; iEGL, high DAPI, low LacZ positive, and Calbindin negative; PCL, low DAPI, LacZ negative, and Calbindin positive; bPCL, medium DAPI, medium LacZ positive, and Calbindin negative; IGL, high DAPI, LacZ and Calbindin negative. P21 and P90 were similar, with three layers including ML (molecular layer), PCL and IGL. N≥3 per group were analyzed for (A-C) (D) X-gal staining of P6 sagittal brain sections of Eu; Gli1-LacZ and Ts65Dn; Gli1-LacZ. Hippocampus and cerebellum areas were compared (n=3 per group). (E) Beta-Glo assay of P6 hippocampus and cerebellum of Eu;Gli1-LacZ and Ts65Dn;Gli1-LacZ (n=4 per group). (F) Beta-Glo assay of P6 hippocampus from 5 pairs of “TRE-hShh;Gli1-LacZ and Camk2a-hShh;Gli1-LacZ” littermates and P6 cerebellum from 5 pairs of “TRE-hShh;Gli1-LacZ and Pcp2-hShh;Gli1-LacZ” littermates. Data were analyzed by paired t-test. Data were represented as mean ± SEM. Data were analyzed by two-way ANOVA and Sidak’s multiple comparisons (E), and by paired t-tests (F).

X-gal staining showed that Ts65Dn;Gli1-LacZ mice had reduced LacZ levels in both hippocampus and cerebellum at P6 compared to Eu;Gli1-LacZ (**Figure 7D**).Quantitative measurements of LacZ levels by Beta-Glo assay showed that trisomy decreased LacZ activity by ∼40% in hippocampus and by ∼20% in cerebellum at P6 (**Figure 7E**).

To test whether Camk2a-hShh and/or Pcp2-hShh enhanced the Gli1 activity in P6 hippocampus and cerebellum, triple transgenic mice, Camk2a-tTA;TRE-hShh;Gli1-LacZ (Camk2a-hShh;Gli1-LacZ) and Pcp2-tTA;TRE-hShh;Gli1-LacZ (Pcp2-hShh;Gli1-LacZ), were generated. Compared to the control (TRE-hShh;Gli1-LacZ), Camk2a-hShh;Gli1-LacZ increased LacZ activity by ∼25% in hippocampus, and Pcp2-hShh;Gli1-LacZ increased LacZ activity by ∼10% in cerebellum (**Figure 7F**).

## Discussion

Shh protein as morphogen plays important roles in development and maintenance of many organs/tissues. Previous conditional Shh knock-in mouse models including “(tetO)7CMV-rShh” (Miller et al., 2004) and “CMV-lox-EGFP-STOP-lox-mShh” (Wang et al., 2010) were generated through random integration. The transgene status such as the inserting location and copy number is important, as random integration caused endogenous gene mutations and transgene copy number variation affect phenotypes of transgenic mice. Here, through the TARGATT site-specific transgenesis we generate a new inducible Shh mouse, TRE-hShh, which has one copy of TRE-hShh transgene inserted in the neutral Rosa26 locus of chromosome 6. TRE-hShh has little or no leaky expression in the absence of tTA, and inducibly expresses the dually lipidated hShh-Np. The bidirectional TRE promoter provides trackable Zsgreen1 expression in hShh-expressing cells. By using a hShh transgene, endogenous and transgene expression can be distinguished, thus we are able to distinguish the effects from endogenous or transgene expression and to tell if they affect the expression of each other. Both human and mouse Shh-Np contain 174 amino acids, the only amino acid difference between the two species is that S44 of human Shh-N becomes T44 in mouse Shh-N (Figure S2C). As serine could often be replaced by threonine without affecting protein function and *vice versa*, it is fair to say that human Shh-Np and mouse Shh-Np are functionally identical and thus have the same dose-response from shh-responding cells *in vivo*. With available tissue specific rtTA or tTA driver lines, the TRE-hShh mouse could be used to study Shh function and its therapeutic potential in many different tissues, and our findings and those of others (Badea et al., 2009; Zhu et al., 2007; Zou et al., 2009) suggest that the tTA is likely more efficient than rtTA to drive TRE-transgene expression in brain.

Attention deficit hyperactivity disorder (ADHD), causing behaviors of inattention or hyperactivity/impulsivity or both, affects 8.4% of all U.S. children 2–17 years of age (Danielson et al., 2018). Two pagiopulation-based studies show that the prevalence of ADHD is significantly elevated in children with DS (Ekstein et al., 2011; Oxelgren et al., 2017). Hyperactivity in Ts65Dn has been consistently observed (Escorihuela et al., 1995; Faizi et al., 2011; Fortress et al., 2015). Children with ADHD reportedly have smaller and abnormally shaped basal ganglion including caudate, putamen, and globus pallidus (Halperin and Schulz, 2006; Qiu et al., 2009). The morphology and plasticity of cholinergic interneurons in basal ganglia of Ts65Dn are affected (Di Filippo et al., 2010; Powers et al., 2016), and trisomy 16 embryos shows hypocellularity in basal forebrain (Sweeney et al., 1989). Shh signaling is highly active in developing ganglionic eminence where interneurons are produced and migrate to basal ganglia (DeBoer and Anderson, 2017). Here, we show that Camk2a-hShh but not Pcp2-hShh normalizes hyperactivity in Ts65Dn, consistent with expression of Camk2a-hShh but not Pcp2-hShh in basal ganglia (Figure 2E VS Figure 3C). Our findings support future study on whether Shh response deficit exists in trisomic basal ganglia cells and if it leads to hyperactivity in Ts65Dn. To validate Shh therapeutic targeting basal ganglia in people with DS or ADHD, more evidence about the relationship between basal ganglia abnormalities, hyperactivity severities, and levels of Shh response deficits are to be discovered.

Improving cognitive function is a major goal to improve life opportunities for people with DS. Shh overexpression in forebrain but not cerebellum significantly improved spatial learning and memory in both 3-month-old Eu and Ts65Dn mice. Further, ageing trisomic mice that expressed hShh throughout life did not show the same loss of learning ability seen in ageing Ts65Dn. Deficits in spatial memory of Ts65Dn are not completely rescued by forebrain Shh overexpression, but the improvement is very significant. Our results indicate that the reduced response to Shh in forebrain rather than cerebellum is the dominant contributor to spatial memory deficits in Ts65Dn.

Gorlin syndrome (GS) is caused by loss-of-function mutations in *PTCH1*, resulting in upregulated Shh signaling, and 3-5% of people with GS develop childhood medulloblastoma (Vorechovsky et al., 1997). Mice heterozygous for *Ptch1* develop medulloblastoma beginning at an early age and 14% are affected by 10 months of age (Wetmore et al., 2000). Sporadic medulloblastoma sometimes presents with mutations in *PTCH1, SUFU,* and *SMO* genes, as well (Dellovade et al., 2006). However, all these mutations are in Shh-responding cells, causing constitutively active Shh signaling in the absence of Shh ligand. Up-regulation of Shh signaling by SAG shows therapeutic efficacy in various animal models such as rats with spinal cord injury (Bambakidis et al., 2010), mice with glucocorticoid-induced neonatal cerebellar injury (Heine et al., 2011), and Ts65Dn mice (Das et al., 2013), and none of these studies show that SAG promotes tumor formation. Mouse Shh ligand overexpression in the mature exocrine pancreas by Ela-CreERT2;LSL-mShh or Ela-CreERT2;LSL-mShh;LSL-mSmo for up to 12 months is insufficient to induce neoplasia (Fendrich et al., 2011). Here, overexpressing hShh ligand in forebrain of Eu and Ts65Dn by Camk2a-hShh for 600 days or in cerebellum by Pcp2-hShh for 14 months does not affect the survival rate. Thus, it is possible that in the absence of underlying mutations in Shh-responding cells up-regulation of Shh signaling by Shh ligand or by SAG is insufficient to induce carcinogenesis. Nevertheless, of the role of Shh signaling in various cancers merits attention in translating Shh research into clinical application.

We show that Gli1-LacZ labels neural precursors in neurogenic niches such as hippocampal DG and cerebellar oEGL, and that P6 Ts65Dn;Gli1-LacZ has reduced LacZ levels in both forebrain and cerebellum comparing with Eu;Gli1-LacZ. As it can take over 2 weeks for LacZ protein to be fully degraded, the Gli1-LacZ reduction in Ts65Dn;Gli1-LacZ is the accumulated readout from P6 to an earlier day. Interestingly, the 20% reduction of LacZ activity in cerebellum of P6 Ts65Dn;Gli1-LacZ is close to the 21% reduction in mitotic GCP of P0 Ts65Dn and the 28% reduction in total GCP of P6 Ts65Dn (Roper et al., 2006). Our findings suggest that Ts65Dn may have neurogenic deficiency in the forebrain and cerebellum during the perinatal and early postnatal development, which is associated with Shh signaling deficiency in these regions. However, whether people with DS have deficits in Shh signaling dependent neurogenesis needs to be further studied, and more research on efficacy, toxicity, and formulation of potential Shh therapy are also required.

## Supporting information

key sources table

## Acknowledgments

We would like to acknowledge Karen Hazzard (NIH/NHGRI) for providing TRE-LacZ mice, JHU Transgenic Core Laboratory, JHU Microscope Facility (MicFac), and Zhipeng Hou of JHU NMR Service Center. This work was supported by NIH grants R01HD038384 (to R.H.R.), R21HD098540-01 (to R.H.R.). The conclusions presented here are not necessarily those of the National Institutes of Health.

## Author Contributions

F.J.G. and R.H.R. conceived the research project. F.J.G. and R.H.R. directed the study. F.J.G. and D.K. performed the experiments. F.J.G., D.K., and B.D. aided with the TRE-hShh transgenic mouse generation and the mouse colony management. B.C. aided with FACS experiments and performed the mass spectrometry analysis. A.S. aided with behavioral tests. D.W aided with MRI analysis. Y.L. M.G.P provided general support. F.J.G. and R.H.R. wrote the manuscript.

## STAR Methods

### Resource Availability

#### Lead Contact

Further information and requests for resources and reagents should be directed to and will be fulfilled by the Lead Contact, Roger H. Reeves

#### Materials Availability

All requests for resources and reagents should be directed to and will be fulfilled by the Lead Contact author. Materials will be made publicly available either through publicly available repositories or via the authors upon execution of a Material Transfer Agreement.

#### Data and code availability

All data are available in this paper or uploaded as supplemental information.

### EXPERIMENTAL MODEL AND SUBJECT DETAILS

#### Animal research

This study was carried out in accordance with the recommendations of the NIH Guide for the Care and Use of Laboratory Animals and the Johns Hopkins University (JHU) Institute of Animal Care and Use Committee. The protocol was approved by the Johns Hopkins University Institute of Animal Care and Use Committee. Mice were maintained in a JHU animal facility with 14-hour light/10-hour dark cycle, temperatures of 65-75°F (∼18-23°C) with 40-60% humidity, and feed with standard chow and in-cage automatic water unless otherwise stated. Mice models including Camk2a-rtTA, Pcp2-rtTA, Camk2a-tTA, Pcp2-tTA, TRE-hShh, Ts65Dn, and Gli1-LacZ, had been backcrossed into C57BL/6J background for more than 6 generations in our lab before any biochemical analysis or behavioral test. For TRE-LacZ mice, homozygous embryos (PMID 7937760, JAX 002621) were gifted from Hazzard, Karen (NIH/NHGRI) and recovered by JHU Transgenic Core Laboratory and maintained as a homozygous strain in SJL/J background throughout the project. The detailed information of genetic background and references was shown in key sources table. The mouse colony management was assisted by Softmouse (Iseehear).

For animal research, we followed the ARRIVE guidance (https://arriveguidelines.org/) to design and carryout the projects. All experimental mice were generated through natural mattings, and a consolidated table of demographic animal information for each experiment of each figure, which included genetic background, age, gender, and animal number, was provided (**Table S2**). We further reiterated this animal information in each figure and/or legend. For biochemical analysis of animal tissues, mice/tissues were appropriately genotyped and assigned with unique identifications (IDs), and investigators were blind to sample genotypes when data were generated. For behavioral tests, each mouse had a unique ID, and each cohort of mice were randomized after being genotyped. Investigators of behavior tests were blind to mouse genotypes. The details of animal generation and experimental procedures were described in the result section and STAR Methods.

For genotyping, a mouse tail tip was placed in a 1.5 ml tube containing 600 ul of lysis buffer (50mM Tris pH8, 100 mM EDTA, 0.5% SDS, and 400 mM NaCl) with 15 ul of 20 mg/ml Proteinase K (ThermoFisher) and incubating at 55°C overnight. The tube was added 180 ul the saturated NaCl solution, and mixed well and then centrifuged at 13,000 rpm for 10 min at 4°C. The supernatant was transferred to another 1.5 ml tube, and filled with 700 ul 100% EtOH, and gently mixed to precipitate DNA and then centrifuged at 13,000 rpm for 10 min at 4°C. The DNA pellet was mixed with 500ul 70% EtOH and centrifuged at 13,000 rpm for 10 min at 4°C, and dried and resuspended in 500 ul DEPC treated H_2_O. For TRE-LacZ genotyping, the primer sets “TRE-LacZ F1/TRE-LacZ R1” and “1179 bGal/bGal 4655” were used. For Camk2a-rtTA genotyping, the primer sets “Camk2a-rtTA F/ Camk2a-rtTA R” and “rtTA and tTA F/ rtTA and tTA R” were used. For Pcp2-rtTA genotyping, the primer sets “Pcp2-rtTA F/ Pcp2-rtTA R” and “rtTA and tTA F/ rtTA and tTA R” were used. For Camk2a-tTA genotyping, the primer sets “Cam2a-tTA F/ Cam2a-tTA R” and “rtTA and tTA F/ rtTA and tTA R” were used. For Pcp2-tTA genotyping, the primer sets “Pcp2-tTA F/ Pcp2-tTA R” and “rtTA and tTA F/ rtTA and tTA R” were used. For TRE-hShh genotyping, the primer sets “ZS306 F/ ZS306 R” and “TRE3G F/ TRE3G R” were used, and homozygous TRE-hShh mice were identified with negative PCR using the “SSL-Rosa11/Rosa10” primer sets. For Ts65Dn genotyping, the primer sets “C17F/ C16R” and “IMR5/ IMR6” were used. For Gli1-LacZ genotyping, the primer sets “Gli1-lacZ Common F/ Gli1-lacZ Mutant R/ Gli1-lacZ WT R” were used. All primer sequence information was shown (**Table S3**).

#### Construction of the targeting vector, pBT378-TRE-bi-hShh-Zsgreen1

The full length hShh cDNA was amplified from HsCD00082632 (DNASU) with primers of SHH-halfRV and SHH-Bam2 and digested with BamHI and then ligated with pTRE3G-bi-ZsGreen1 (TaKaRa) after BamHI/EcoRV digestion to generate the first intermediate plasmid, pTRE3G-bi-hShh-Zsgreen1. The pTRE3G-bi-hSHH-Zsgreen1 vector was digested with PciI and filled-in using Klenow followed by EcoRI digestion to create the TRE-hShh cassette, which was ligated with pBS-SK(+) after SmaI/EcoRI digestion to create pBS-SK(+)-TRE-hShh. Zsgreen1/polyA, amplified from pTRE3G-bi-ZsGreen1 with primers of ClaI-F_Zs2 and RI-R_Zs, was ligated with pBS-SK(+)-TRE-hShh after ClaI/EcoRI digestion to generate the second intermediate plasmid, pBS-SK(+)-TRE-bi-hShh-Zsgreen1. The pBS-SK(+)-TRE-bi-hShh-Zsgreen1 vetetor was digested with ClaI and Not I to get the fragment of TRE-bi-hShh-Zsgreen1 with polyAs, which was ligated with pBT378 after ClaI/Not I digestion to generate the targeting vector, pBT378-TRE-bi-hShh-Zsgreen1 (also known as the TRE-hShh targeting vector, **Figure S1A**).

#### Generation of TRE-hShh mice using TARGATT™ site-specific knockin technology

In JHU Transgenic Core Laboratory, 50ng/ul φC31mRNA and 3ng/ul TRE-hShh targeting vector were diluted in RNAse free injection buffer (10 mM Tris-HCl, pH 7.4, 0.25 mM EDTA) were injected into the pronucleus of 297 1-cell embryos from Rosa26 TARGATT mice (Applied StemCell), which were transferred into the oviducts of 11 pseudopregnant ICR moms. Founders were identified by PCR of tail DNA using primer sets SSL and SSR (Applied StemCell) that were specific for the right and left junctions of the attP/Rosa26 locus. The first round of injection yielded 0 founder from 16 pups with the transgene of interest, and the second round of injection yielded 3 founders from 15 pups. Two lines had identical attP1/attP3 insertions, and the other one had only attP3 insertion.

To verify transgene integration, Southern blot analysis was used. The 5’- and 3’-probes were made by PCR with primer sets Tsh5prF/Tsh5prR of WT genomic DNA and Tsh3prF2/Tsh3prR3 of pROSA26-PA (plasmid #21271, Addgene), respectively, and then labeled with α-32P-dCTP (Amersham Rediprime II Random Prime Labelling System). Genomic DNA from 3 founder lines and wild type were digested with HindIII, separated on 1x TBE 0.9% agarose gel at 50 V overnight with recirculation and transferred to Hybond N+, which was hybridized with 5’- or 3’-probes. The membrane was imaged on Molecular Dynamics Storm Imager and then exposed to film.

#### Cell culture and transfection

To generate MEF, E14.5 mouse embryos were transferred into 50ml tube with 30ml sterile PBS. Heads and liver/organs were removed with sterile razor blade. The remaining tissue were rinsed with PBS and placed in one well of a 6 well plate and then dissociated into fine pieces and digested in 3ml of 0.25% trypsin-EDTA at 37°C for 20 min. The cell solutions were then transferred into a 10cm plate containing 10 ml of MEF culture medium (DMEM, 10% FBS, 0.1 mM β-mercaptoethanol, 50 U penicillin, and 50 μg/ml streptomycin) and transferred to 37°C incubator overnight. In the next day, the culture plates were replaced with 12ml fresh MEF culture medium and replaced with fresh medium every 2 days thereafter. To test the targeting vector, MEF cells were co-transfected with CMV-rtTA (pCMV-Tet3G, Clontech) and the TRE-hShh targeting vector using Lipofectamine 3000 (ThermoFisher). After 16 hours, Doxycycline (Dox, Sigma-Aldrich) was added at 0, 0.1,1 ug/ml for 32 hours. Cells were analyzed by Taqman RT-PCR, western blot, and immunostaining.

#### Doxycycline delivery to mice

To test Camk2a-rtTA and Pcp2-rtTA mice, Camk2a-rtTA;TRE-LacZ and Pcp2-rtTA;TRE-LacZ were generated through crossing TRE-LacZ with Camk2a-tTA and Pcp2-tTA, respectively. Beside 625 mg/kg food diets (Envigo), the Dox (Sigma-Aldrich) that induced hShh overexpression in CMV-rtTA and TRE-hShh co-transfected MEF cells was used to make 2 mg/ml or 3.5mg/ml Dox drinking water containing 2% sucrose. Mice were treated with different doses and times of Dox (**Table S1**) and stained with X-gal at P30.

#### Gene expression analysis

For Taqman RT-PCR, the total RNAs were extracted by Trizol (ThermoFisher) and Chloroform (MilliporeSigma) and followed by the RNeasy column purification (QIAGEN. Cell pellets or homogenized tissues were mixed with 500 ul Trizol/sample (1ml for larger samples) and then 100 µl chloroform (the Trizol/chloroform ratio was 5) and incubating at room temperature for 10 min. The sample was then centrifuged at 13,000 rpm for 20 min at 4°C, and the upper, aqueous phase was transferred to a new tube and mixed with 1 volume of 70% EtOH/DEPC H_2_O. The sample was transferred to the RNeasy column of RNeasy Mini Kit (QIAGEN) and centrifuged at > 9000 rpm for 20 s. The flow-through was discarded and repeated the centrifugation with the remaining sample. The Rneasy column was added with 700 µl Buffer RW1 and centrifuged at > 9000 rpm for 20 s, and flow-through was discarded. The RNeasy column was added with 500 µl Buffer RPE and centrifuged at > 9000 rpm for 20 s, and flow-through was discarded. The RNeasy column was added with another 500 µl Buffer RPE and centrifuged at > 9000 rpm for 2 min, and flow-through was discarded. The column was transferred into a new 2 ml collection tube and centrifuged at full speed for 1 min. The column was transferred to a new (labelled) 1.5 ml tube and added with 30-50 µl RNase-free water and sitting at room temp for ∼2 min, then centrifuged for 1 min at > 9000 rpm. The flow-through (RNA samples) was quantified using nanodrop and stored at −80°C. The cDNAs were synthesized using high-capacity cDNA reverse transcription kits according to the manual (ThermoFisher). The TaqMan Gene Expression Master Mix (ThermoFisher) and Taqman probes were used to perform quantitative RT-PCR using QuantStudio 6 Flex Real-Time PCR Systems (ThermoFisher).

For Western blot, cell pellets or homogenized tissues were lysed in 500ul NP-40 lysis buffer (50 mM Tris (pH 7.5), 0.1% NP-40, 100mM NaCl, 1 mM MgCl2, 5 mM EDTA) supplemented with 1X protease inhibitor cocktail and 1X Halt phosphatase inhibitor cocktail on ice for 30 min. Cell lysates were sonicated for 10 pulses at level 1 with 10% output 3 times, and then incubating on ice for another 20 min. Soluble lysates were obtained by centrifugation at 17,000× g for 20 min at 4°C, and the protein concentration was measured by BCA (ThermoFisher). Protein extracts were separated by SDS-PAGE, and transferred to Nitrocellulose membranes. The Nitrocellulose membranes were probed with primary and secondary antibodies that described in key sources table, and then were washed extensively with PBS-T (0.1% Tween-20 in PBS) after each antibody probe.

For immunostaining cells, cells were growing and transfected on poly-D lysine coated coverslips, and then fixed in 4% PFA for 15 min. The cells were incubated with blocking buffer (5% normal goat serum and 0.1% triton X-100) for 1 hour at room temperature, and primary antibodies at 4°C overnight (or room temperature for 2 hours), and then Alexa Fluor conjugated secondary antibodies (ThermoFisher) for 45-60 min and then incubated in DAPI solution (1ul original DAPI diluted in 50ml PBS-T) for 10 min.

Coverslips were mounted on slides with “ProLong Gold Antifade Mountant with DAPI” (ThermoFisher) and dried, and then imaged using Zeiss LSM800 GaAsP (Microscope Facility, Johns Hopkins School of Medicine).

#### Histology

For Xgal staining of brain sections, mice were perfused with PBS and 4% PFA. Brains were isolated and kept in 4% PFA overnight and embedded into optimal cutting temperature compound (OCT compound) and were sectioned at 30 um using Leica Cryostat CM 3050S. The sections were mounted on the glass slides and dried by natural air. Slides were fixed in fixation buffer (2% paraformaldehyde, 0.02% glutaraldehyde, 2 mM MgCl2 in PBS) for 10 min at room temperature, and washed in PBS for 10 min, and then incubated in washed solution (5 mM EGTA, 0.01% Deoxycholate, 0.02% NP40, 2 mM MgCl2 in 0.1M phosphate buffer) for 10 min, and then immersed in X-gal staining solution (5 mM K_3_Fe(CN)_6_, 5 mM K_4_Fe(CN)_6_, 5 mM EGTA, 0.01% Deoxycholate, 0.02% NP40, 2 mM MgC12, 1mg/ml X-gal) overnight at 37°C in the dark. Slides were rinsed in PBS for 10 min and washed in distilled water for 5 min. Some slides were counter-stained with nuclear fast red for 1 min, rinsed in distilled water once, and washed in distilled water for 1 min, dehydrated, and mounted with DPX and cover glasses.

For Immunostaining for brain sections, frozen sections (30 or 40 um) were fixed in 4% PFA for 30 min and rinsed twice with PBS. The sections were permeabilized and blocked with blocking buffer (0.5% triton X-100 and 10% goat serum in PBS) for 1 hour. The sections were probed with primary antibodies diluted in blocking buffer at 4°C overnight or at room temperature for 2 h (negative control was not treated with primary antibodies), followed by 3 washes in PBS-T for 10 min each, and then stained with Alexa Fluor conjugated secondary antibodies diluted in blocking buffer for 1 h followed by 3 washes in PBS-T, 10 min each. The sections were incubated in DAPI solution (1ul DAPI diluted in 50ml PBS-T) for 10 min, and then mounted on slides with “ProLong Gold Antifade Mountant with DAPI”. The Slides were kept in dark overnight at room temperature and imaged using Zeiss LSM800 GaAsP. To create movies from confocal images, Z-stack and tile scan images were stitched in Zen (Zeiss) and converted to a movie in Imaris (Bitplane).

#### Flow cytometry

Mice (TRE-hShh, Camk2a-tTA;TRE-hShh, and Pcp2-tTA;TRE-hShh) were euthanized with isoflurane and perfused with cold PBS. Cerebellum, hippocampus, and cerebral cortex were dissected, and then cells were dissociated using Papain Dissociation System (Worthington-biochem). Tissues were first incubated with a proper amount of the papain/DNase solution in 15ml tube at 37°C for 10 min. The tissues were then triturated by a 10 mL pipette for 10 times, and by a 5ml pipette for 10 times, and by a 1ml pipette tips for 15 times. The tubes were placed on ice for 10 min to allow any remaining chunks of tissues to settle to the bottom, and the supernatants were transferred into a new 15 mL tube, followed by 2000 rpm centrifugation at 4°C for 5 min. Cell pellets were resuspended in re-suspension media (2.7 mL of EBSS were added with 300 μL albumin-ovomucoid inhibitor and 150 μL DNase). The cell suspension was passing through a 70 μm cell strainer (MilliporeSigma) to remove tissue debris, and then carefully loaded on the top of 5 mL albumin inhibitor solution in 15 min tube to create a continuous density gradient and centrifuged at 1000 rpm for 5 minutes at 4°C. The cell pellets were resuspended in re-suspension media and passing through 40 μm cell strainers (BD Biosciences) into a new 15 mL tubes on ice.

The levels of Zsgreen1 expression for transgenic mice were analyzed in SH800 Cell Sorter (Sony Biotechnology Inc, San Jose, CA). Cells were with illuminated with a 488-nm laser and the fluorescence was determined using the FL1 525 ± 50 nm emission filter. The minimal cell count of each sample was 200,000. In the FlowJo software v.10.1r7 (Ashland, OR), the same GFP-positive gating was used for each group to differentiate GFP positive and negative cells. The percentage of GFP positive (GFP+) was calculated by the total number of GFP positive cells divided by the total cell count.

#### MALDI-TOF MS analysis

The purity and concentration of recombinant Shh-N protein including recombinant mouse Shh-N (Cys25-Gly198, with a C-terminal 6-His tag, R&D 461SH), human Shh-N (an N-terminal Ile-Val-Ile sequence substituted for the naturally occurring Cys25 residue, PeproTECH 100-45), and human Shh-Np (R&D 8908-SH/CF) was first confirmed by Coomassie blue staining and Western blot (**Figure S2B**). To make the matrix solution, 10 mg sinapinic acid was dissolved in 1 mL of 50:50 water/acetonitrile containing 0.1% trifluoracetic acid. 1 µL matrix was added on the MTP 384 target plate (Bruker) and dried, followed by depositing 1 µL protein, and then added another 1 µL matrix to the plate and dried. The MALDI-TOF spectra were acquired on a Bruker AutoFlex III (Billerica, MA) using positive ion mode.

#### Behavior tests

3-month-old male mice of the forebrain-cohort (Eu;TRE-hShh (n=16), Eu;Camk2a-hShh (n=14), Ts65Dn;TRE-hShh (n=15), and Ts65Dn;Camk2a-hShh (n=14)) and the cerebellum-cohort (Eu;TRE-hShh (n=21), Eu;Pcp2-hShh (n=20), Ts65Dn;TRE-hShh (n=19), and Ts65Dn;Pcp2-hShh (n=18)) were used for behavioral tests in the sequence, open field, visual discrimination water maze test, MWM, and RRWM (**Figure 4A**). The ANY-maze tracking system (Stoelting Co.) was used to collect data.

The open field tests were performed after three days of handling in the same room consisting of indirect diffusing light (∼150 lux). The whole arena size was 37 cm X 37 cm, and the center area (21.6 cm X 21.6 cm) was 34% of the whole arena. Distance traveled and percentage of time spent in center were analyzed. One Ts65Dn;TRE-hShh mouse in the forebrain-cohort and one Eu;Pcp2-hShh of the cerebellum-cohort were excluded from statistical analysis because of bad tracking.

For visual discrimination (VD), all mice were pre-trained to climb and stay on a submerged platform (10 cm X 10 cm) in a small clear water pool (45 cm diameter) for five trials on the day prior to VD tests. In a water tank 126 cm in diameter, non-toxic white tempera paint was used to make the platform invisible. No spatial cue was used, but the location of the platform was made visible by attaching a black extension that was 4 cm above water surface. During VD tests, the platform position started from W and to E and then to S, and two trials were performed in each platform position (**Figure S5A**).

For classic MWM, the platform remained in the same position in “SE quadrant”, with the water temperature at 22 ± 2°C. With the same spatial cues MWM tests were performed for four training days, and each training day had 10 trials that included 8 acquisition trials and 2 probe trials of short-(30 min) and long-delay (24h), and a longest delay probe trial was conducted on the day 7, 72 h after the last trial of the training day 4 (**Figure S6A**). In acquisition trials, the platform was hidden ∼1.8 cm below the water surface and 60 seconds maximum time was allowed for mice to find the platform, and the tester would visually or manually guide it to the platform if a mouse did not find the platform by itself. In the probe trials that were 30-40 seconds, the platform was lowered to a position that mice were not able to climb onto. At the end of probe trials, the collapsed platform was raised to the same position used in the acquisition trial, and then the tester guided the mouse to the platform, which helped mice maintain the same response-reinforcement contingency of the acquisition. The quadrant target area, a circle inscribed in the platform quadrant, covered ∼17% of water maze tank. Trial 5 of Day 1 was a probe trial drill, during which the platform was lowered to a position that mice were not able to climb onto and mice were only allowed to swim for 10 seconds and then a tester raised the platform and guided mice to the platform, and no data from the trial was included for analysis. If a mouse continually failed to follow the tester’s guidance to reach the platform, it was excluded from analysis, and no mice were excluded for this reason.

Following the classic MWM, the forebrain-cohort containing 12 Eu;TRE-hShh, 12 Eu;Camk2a-hShh, 10 Ts65Dn;TRE-hShh, and 10 Ts65Dn;Camk2a-hShh mice were tested in RRWM without changing any spatial cues. RRWM was consisted of two reversal WM tests (**Figure S7A**). In the first reversal WM, the platform was relocated to NW from SE for two training days. In the second reversal WM, the NW platform was relocated to SW for another two training days. Each training day had 10 trials including 8 acquisition trials and 2 probe trials for short delay (30 min) and long delay (24h). Trial 1 of reversal WM day 1 was the same as the 72h delay probe trial in the classic MWM. This cohort was were re-tested in MWM at ∼7-month-old, which had the same protocol as the classic MWM above. The only differences were that the water tank was in a different room and that spatial cues were different.

#### Brain morphometry by 3D T2-weighted MRI

The 14-month-old cerebellum-cohort (n=8 per group) were used for ex vivo MRI. Mice were perfused with 4% PFA after PBS and heads were post-fixed for 1 week, then kept in PBS for 3 days. Heads were stored in Fomblin to prevent dehydration during imaging in an 11.7 Tesla scanner (vertical bore, Bruker Biospin, Billerica, MA). 3D T2-weighted images were acquired on an 11.7 Tesla Bruker scanner (Bruker Biospin, Billerica, MA, USA) with the resolution = 0.08 mm x 0.08 mm x 0.08 mm, which were first aligned to the template image using automated image registration software (Diffeomap, www.mristudio.org) and adjusted to an isotropic resolution of 0.0625 mm × 0.0625 mm × 0.0625mm. The region of interest (ROI) of the whole brain and cerebellum was manually draw, and the whole brain volume and cerebellar volume were calculated for each mouse, and then the ratio of cerebellum to brain from each group was statistically compared (**Table S4**).

#### Beta-Glo assay

P6 Mice (Eu;Gli1-LacZ and Ts65Dn;Gli1-LacZ littermates, or TRE-hShh;Gli1-LacZ and Camk2a-tTA;TRE-LacZ;Gli1-LacZ littermates, or TRE-hShh;Gli1-LacZ and Pcp2-tTA;TRE-LacZ;Gli1-LacZ littermates) were euthanized. Cerebellum, hippocampus, and cerebral cortex were dissected, which were flash frozen in liquid nitrogen and stored in - 80°C. On the day of beta-Glo assay, tissues of the whole cerebellum or hippocampus were incubated with Reporter Lysis Buffer (Promega) and disrupted using TissueLyser LT (QIAGEN) at 4°C. The homogenized tissues were sonicated at 20% output for 10 s, which was repeated for 3 times. The cell lysis was incubating on ice for 20 min, followed by centrifugation at 17,000× g for 20 min at 4°C. The protein concentration of soluble lysate was determined by BCA protein assay, and then all groups were diluted to the same concentration. 2 ul and 5 ul lysates of each sample were mixed well with 100 μl of Beta-Glo Assay Reagent (Promega) in 96-well plates. The samples were incubating for 30 min at room temperature, and then luminescence was measured with a Wallac 1450 MicroBeta (PerkinElmer). Luminescence was normalized by protein concentration (LCPS/ug), and the relative LCPS/ug of littermates were compared and analyzed by paired t-test.

#### Quantification and statistical analysis

We used GraphPad Prism to do all statistical analysis unless otherwise stated. Generally, we used paired or unpaired t tests to compare differences between two groups, and the one-way ANOVA and post-hoc Tukey’s multiple comparisons test to analyze the significance of one independent variable and compare differences between three or more groups. To analyze the significance of two independent variables and compare differences between three or more groups, the two-way ANOVA and post-hoc Tukey’s or Sidak’s multiple comparisons tests were used. Data were represented as mean ± SEM. All significance thresholds were set at p < 0.05 unless otherwise stated. All the detailed statistical analysis for each figure was available in consolidated tables (**Table S5**).

## Supplemental information

**Figure S1.**
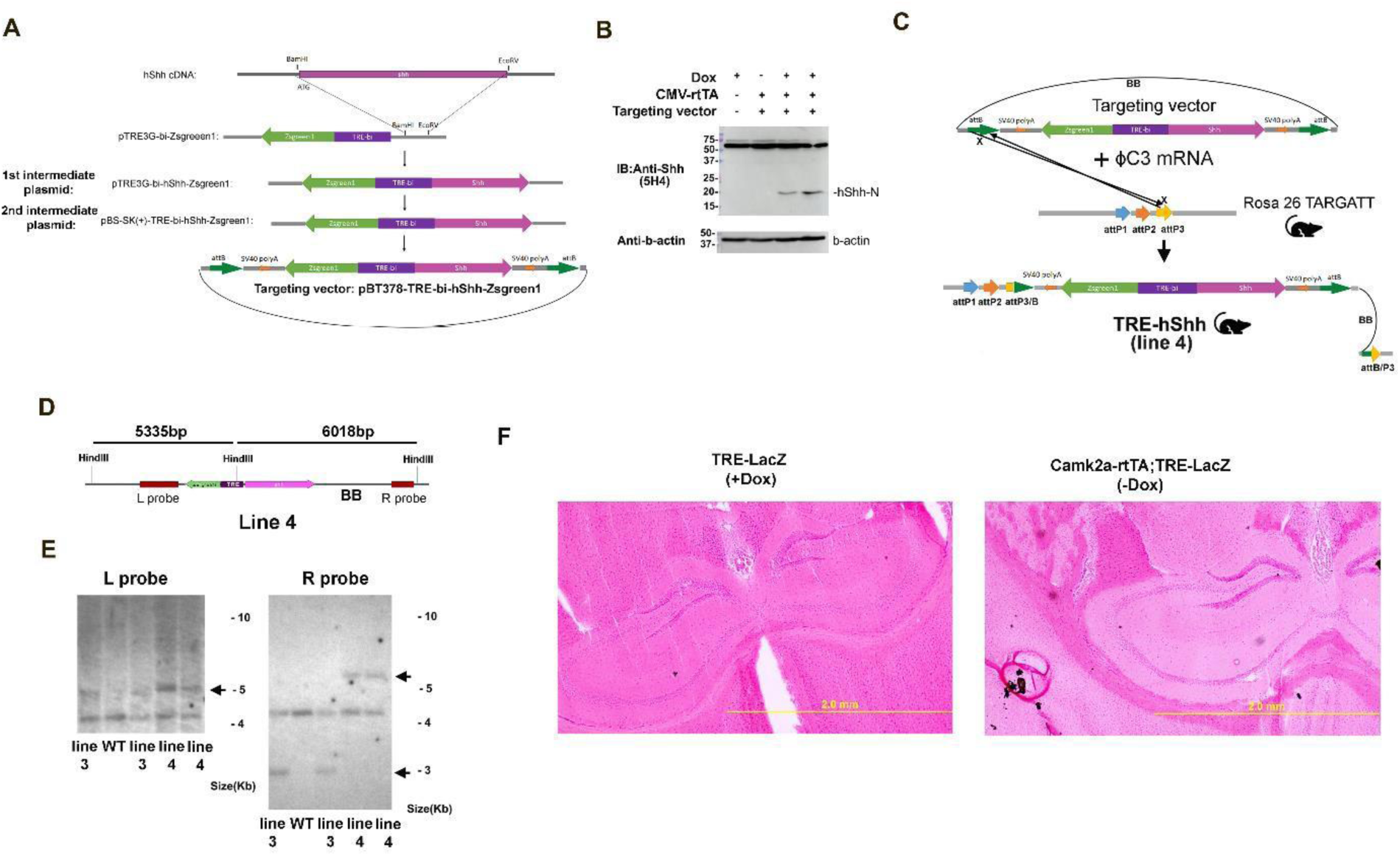
Animal models creation and verification, related to Figure 1. (A) Scheme illustrating the generation of the targeting vector, TRE-bi-hShh-Zsgreen1. For the generation of the first intermediate plasmid, “pTRE3G-bi-hShh-Zsgreen1”, full length hShh cDNA was amplified from HsCD00082632 (DNASU) with primers of SHH-halfRV and SHH-Bam2, which was digested with BamHI and then ligated with pTRE3G-bi-ZsGreen1 after BamHI/EcoRV digestion. To generate the intermediate plasmid, “pBS-SK(+)-TRE-bi-hShh-Zsgreen1”, pTRE3G-bi-hSHH-Zsgreen1 was digested with PciI and filled-in using Klenow followed by EcoRI digestion to create the TRE-hShh cassette, which was ligated with pBS-SK(+) after SmaI/EcoRI digestion to create pBS-SK(+)-TRE-hShh, and Zsgreen1/polyA was amplified from pTRE3G-bi-ZsGreen1 with primers of ClaI-F_Zs2 and RI-R_Zs and ligated with pBS-SK(+)-TRE-hShh after ClaI/EcoRI digestion. For the generation of the Targeting vector, pBS-SK(+)-TRE-bi-hShh-Zsgreen1 was digested with ClaI and Not I to get the fragment of TRE-bi-hShh-Zsgreen1 with polyAs, which was ligated with pBT378 after ClaI/Not I digestion. (B) Western blot of MEFs that were co-transfected with CMV-rtTA and targeting vectors and treated with or without Dox using anti-N-terminus Shh antibodies (5H4). (C) Scheme illustrating that TRE-hShh line 4 had the whole plasmid (with BB) inserted at attP3/attP3. (D) Model of predicting southern blot of TRE-hShh (line4). HindIII digestion produces two fragments. Because of containing extra two attP sites, the left fragment of line 4 was 5335bp slightly larger than 5193bp-fragment of line 2 and 3. Because of containing BB, the right fragment was 6018bp which was significantly larger than 2857bp fragment of line 2 and 3. (E) Southern blots of WT and TRE-hShh (line 3 and 4) mice using radiolabeled L and R probes. (F) Representative images of X-gal stained coronal brain sections from P30 TRE-LacZ mice with Dox treatment from conception (625 mg/Kg food pellets plus 3.5 mg/ml in drinking water) and Camk2a-rtTA;TRE-LacZ without Dox. Hippocampus was shown.

**Figure S2.**
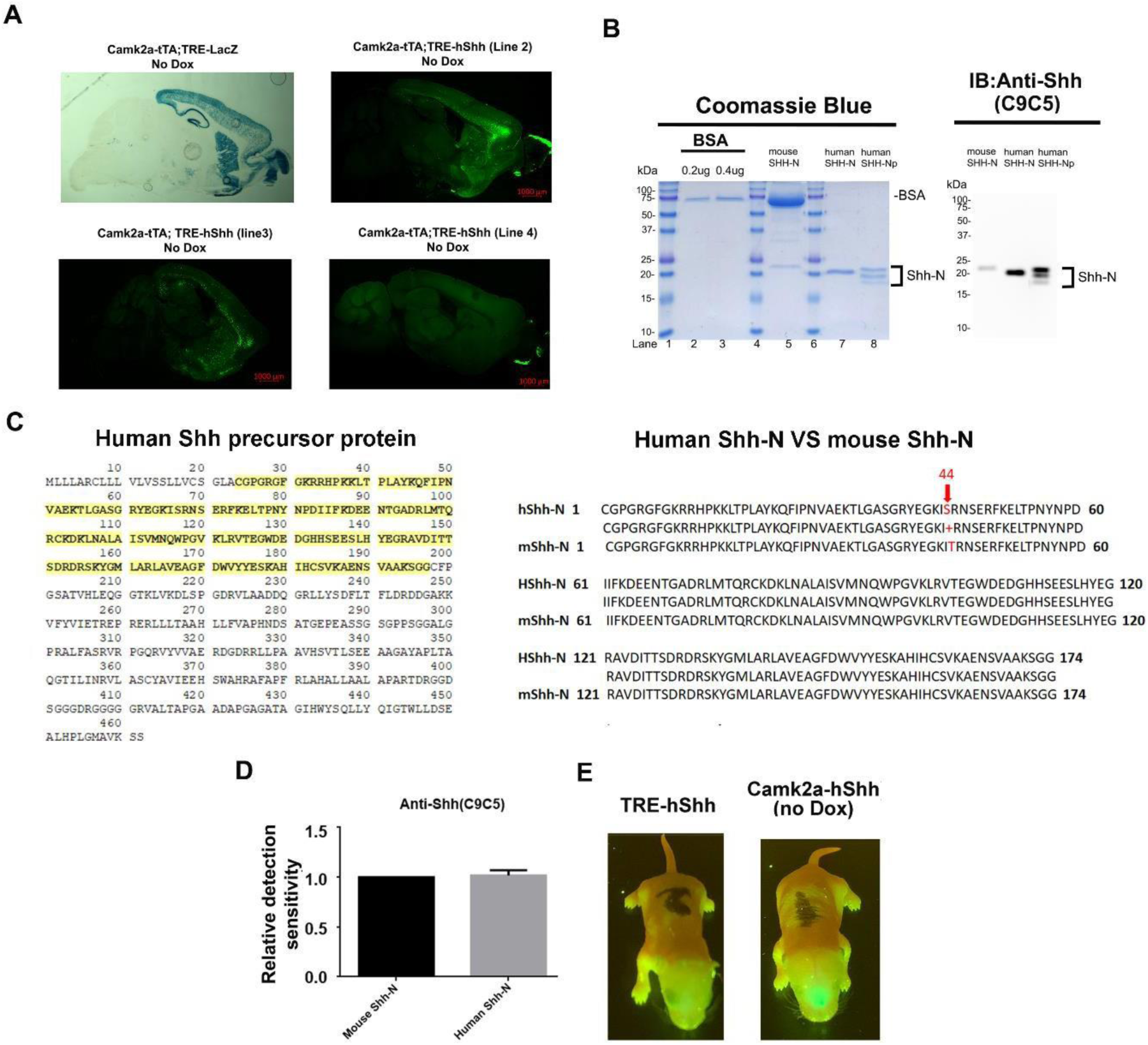
Camk2a-tTA induces TRE-hShh to express hShh in mice, related to Figure 2. (A) Sagittal brain sections of Camk2a-tTA;TRE-hShh (Camk2a-hShh), from lines 2, 3 and 4 mice at 2-month-old that were imaged in Zsgreen1 channel and the tile scan confocal images were shown. Xgal staining of sagittal brain sections of Camk2a-tTA;TRE-LacZ mice were used as the comparison. Scale bar 1000 um. (B) The purified recombinant mouse Shh-N (6-His-tagged, R&D 461SH), human Shh-N (IVI at N-terminus, PeproTECH 100-45), and human Shh-Np (R&D 8908-SH/CF) were shown in Coomassie blue staining (left) and Western blot of anti-Shh (right). The total loaded protein in Coomassie blue staining was 50 times as much as that in Western blot. (C) Full length human Shh protein sequence (left) and the comparison of human Shh-N and mouse Shh-N sequence (right) were shown. The C24-G197 (yellow highlighted) is the Shh-N with MW of 19.56 kDa. Palmitoylation at C24 adds ∼238 Da and cholesterol addition at G197 is ∼369 Da. Thus, Shh-Np with the dual lipidation is ∼20.17 kDa, and with palmitoylation modification only is 19.8 kDa. (D) Quantitative analysis of C9C5’s relative detection sensitivity of mouse Shh-N to human Shh-N. Purified recombinant protein of mouse Shh-N (6-His-tagged, R&D 461SH) and human Shh-N (IVI at N-terminus, PeproTECH 100-45) were compared by Western blot. (E) P1 TRE-hShh and Camk2a-hShh pups without Dox treatment was visualized by GFP flashlight.

**Figure S3.**
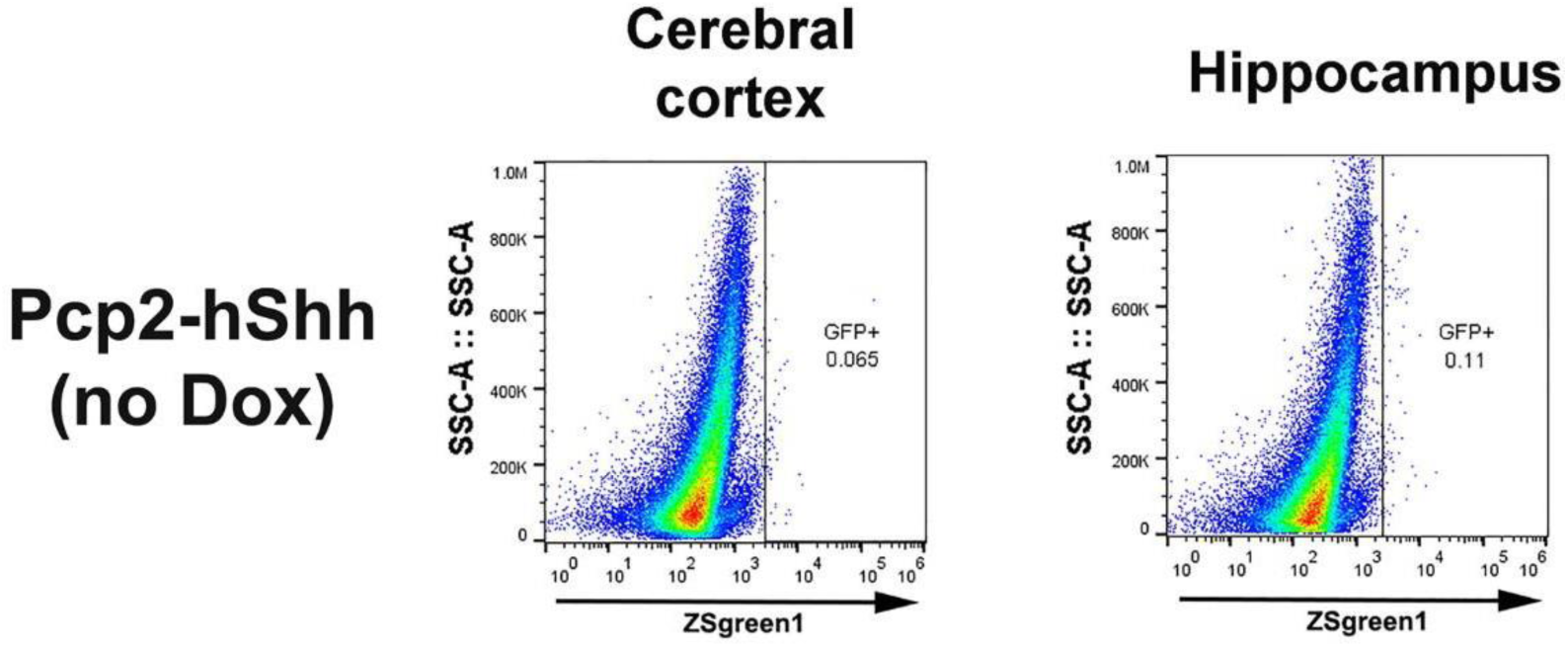
FACS of cerebral cortex and Hippocampus from of Pcp2-hShh without Dox treatment, related to Figure 3.

**Figure S4.**
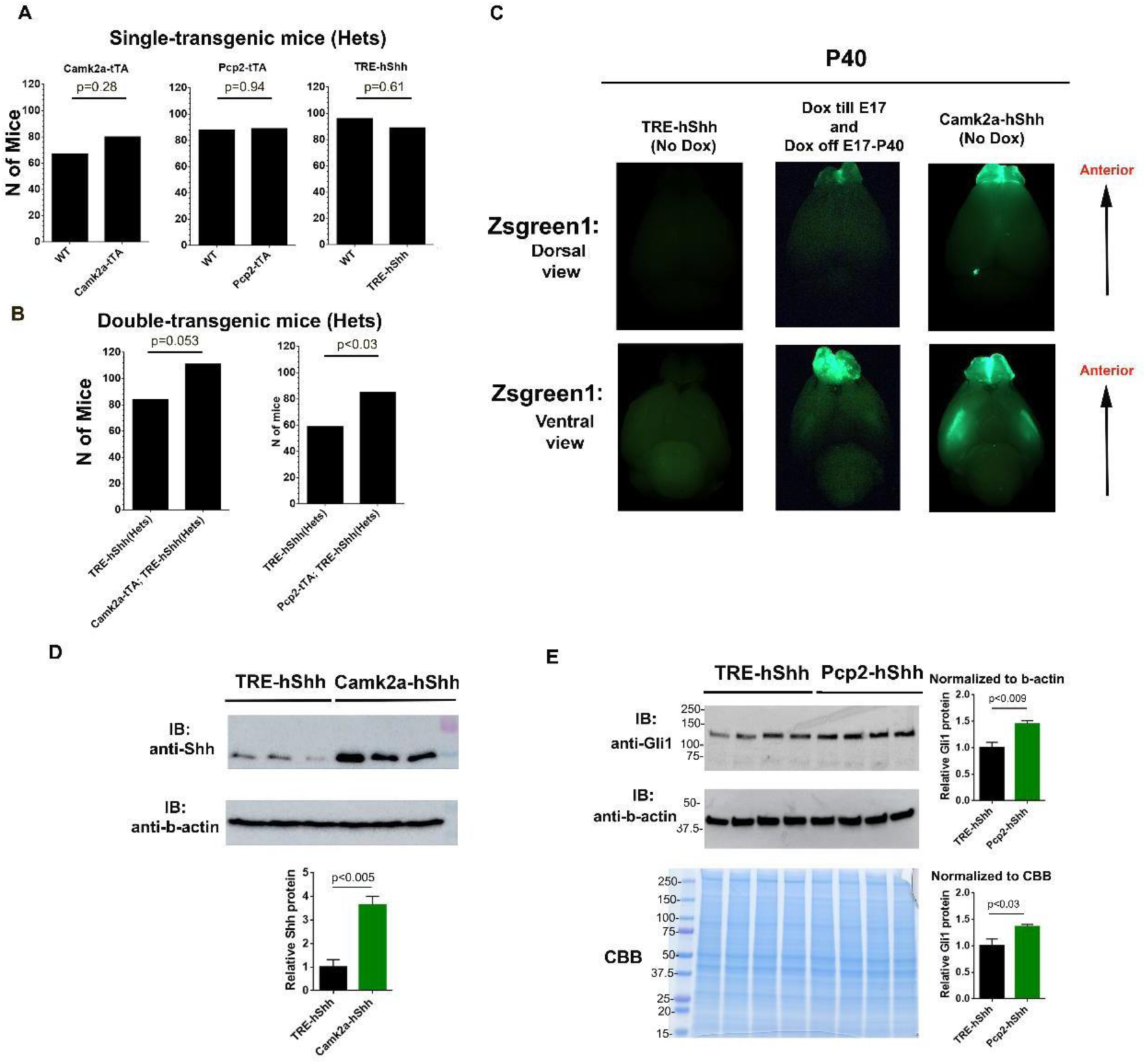
The patterns of transgene inheritance and expression, related to Figure 4. (A) The offspring ratio of single transgenic mice. The breeding records of Camk2a-tTA, Pcp2-tTA, and TRE-hShh single transgenic mice were analyzed at weaning age. Data were analyzed by Binomial test. (B) The offspring ratio of double transgenic mice, Camk2a-tTA;TRE-hShh and Pcp2-tTA-hShh, to TRE-hShh was compared. Data were analyzed by Binomial test. (C) P40 mouse brains of TRE-hShh (left), Camk2a-hShh with “E0-E17” Dox treatment (middle), and Camk2a-hShh without Dox treatment (right), were visualized in GFP-channel. (D) Western blot of cortex from TRE-hShh and Camk2-hShh at 3-month-old, and quantification of Shh density was analyzed by unpaired t-Test (n=3). (E) Western blot of cerebellum from TRE-hShh and Pcp2-hShh mice, and quantification of Gli1 density was analyzed by unpaired t-Test (n=4).

**Figure S5.**
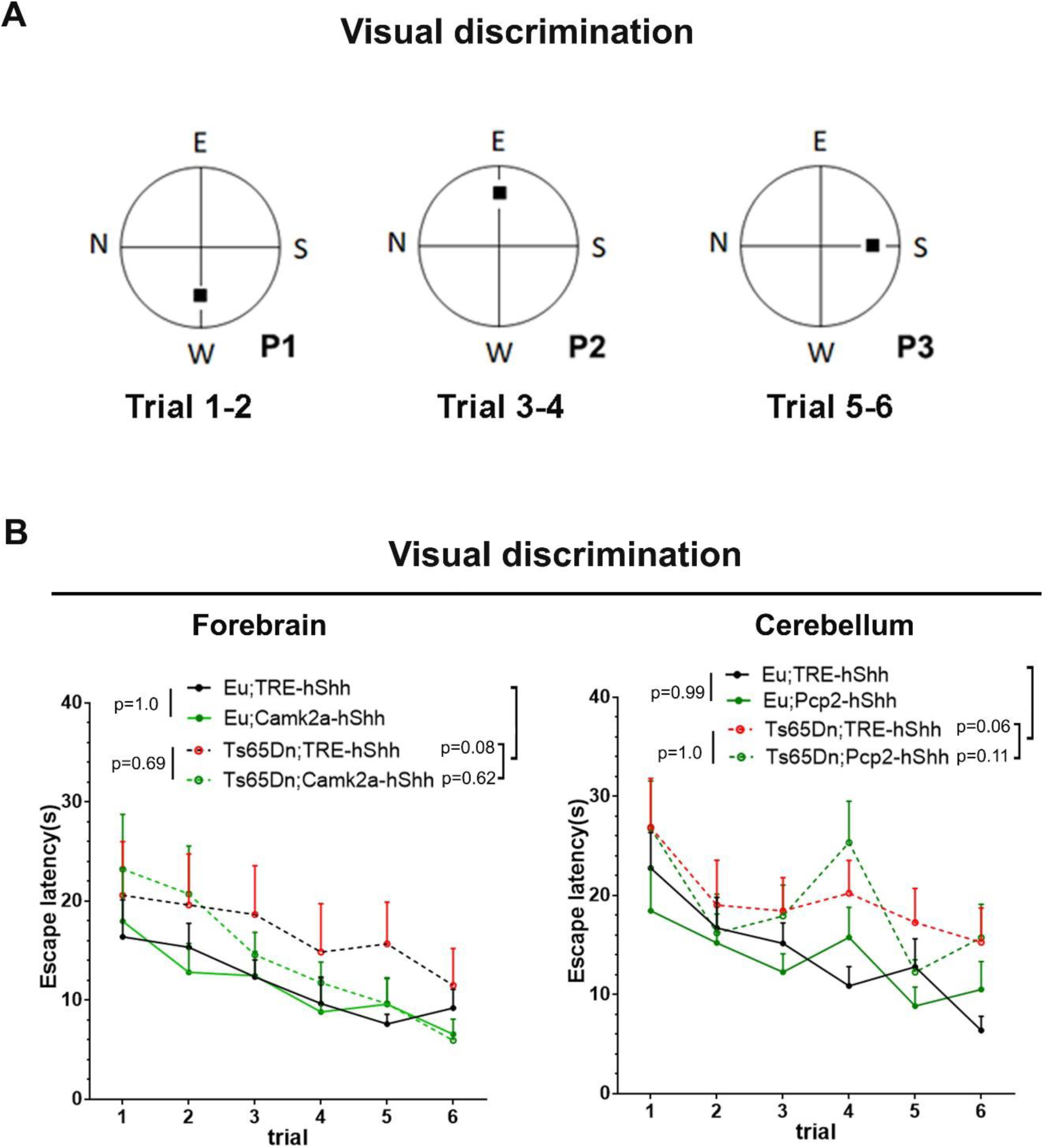
Visual discrimination task, related to Figure 5. (A) Scheme illustrating six-trial visual discrimination task. (B) Visual discrimination task results of the forebrain cohort (left) and the cerebellum cohort (right). Data were represented as mean ± SEM and analyzed by two-way RM AVOVA and Tukey’s multiple comparisons test.

**Figure S6.**
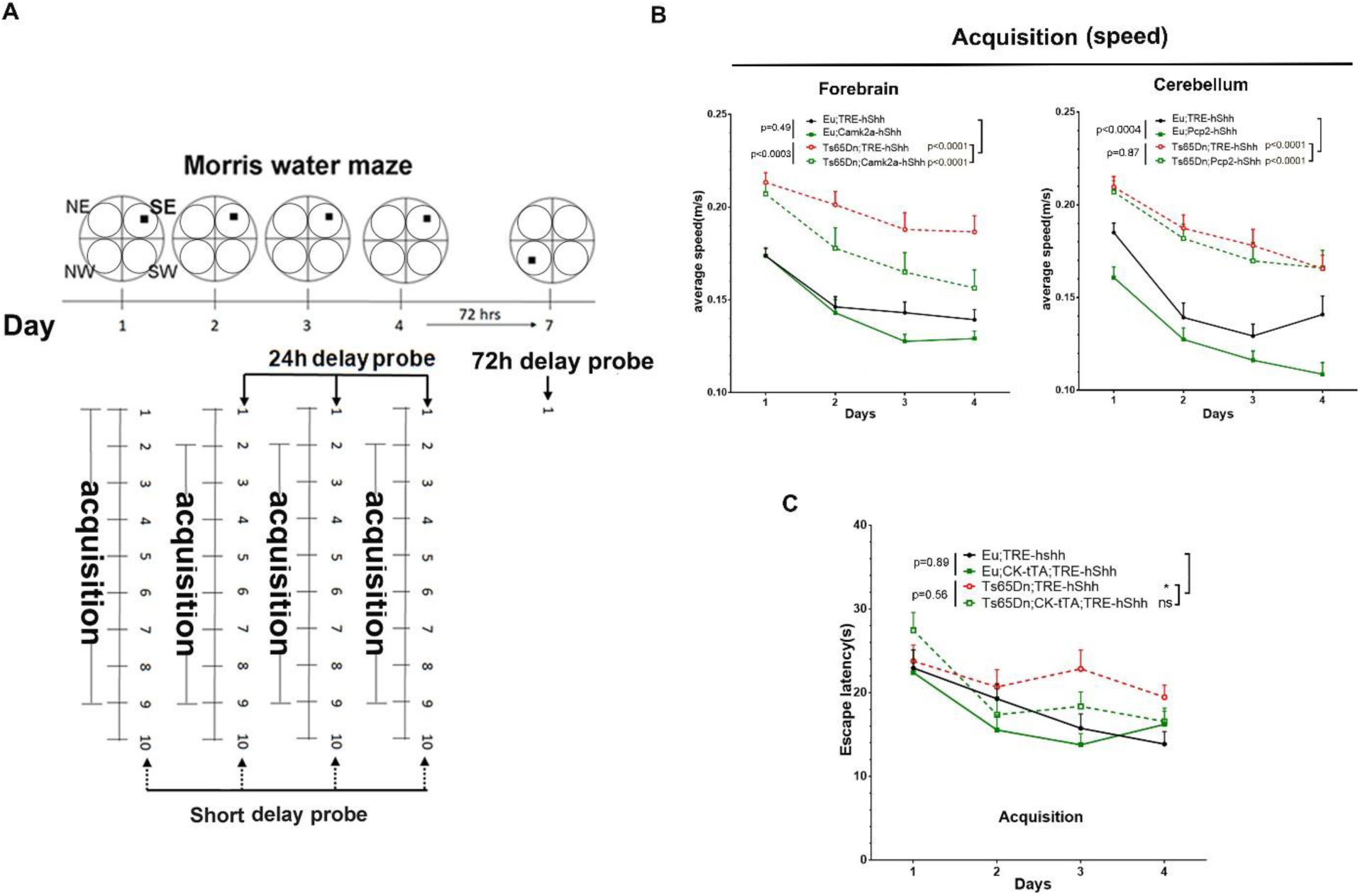
Supplements for MWM, related to Figure 5. (A) Scheme illustrating classic MWM design. (B) Swimming speed of acquisition trials (average speed of 8 trials per day). (C) Escape latency in acquisition trials of MWM (average latency of 8 trials each day). Data are analyzed by two-way ANOVA and Tukey’s multiple comparisons test and expressed as mean ± SEM.

**Figure S7.**
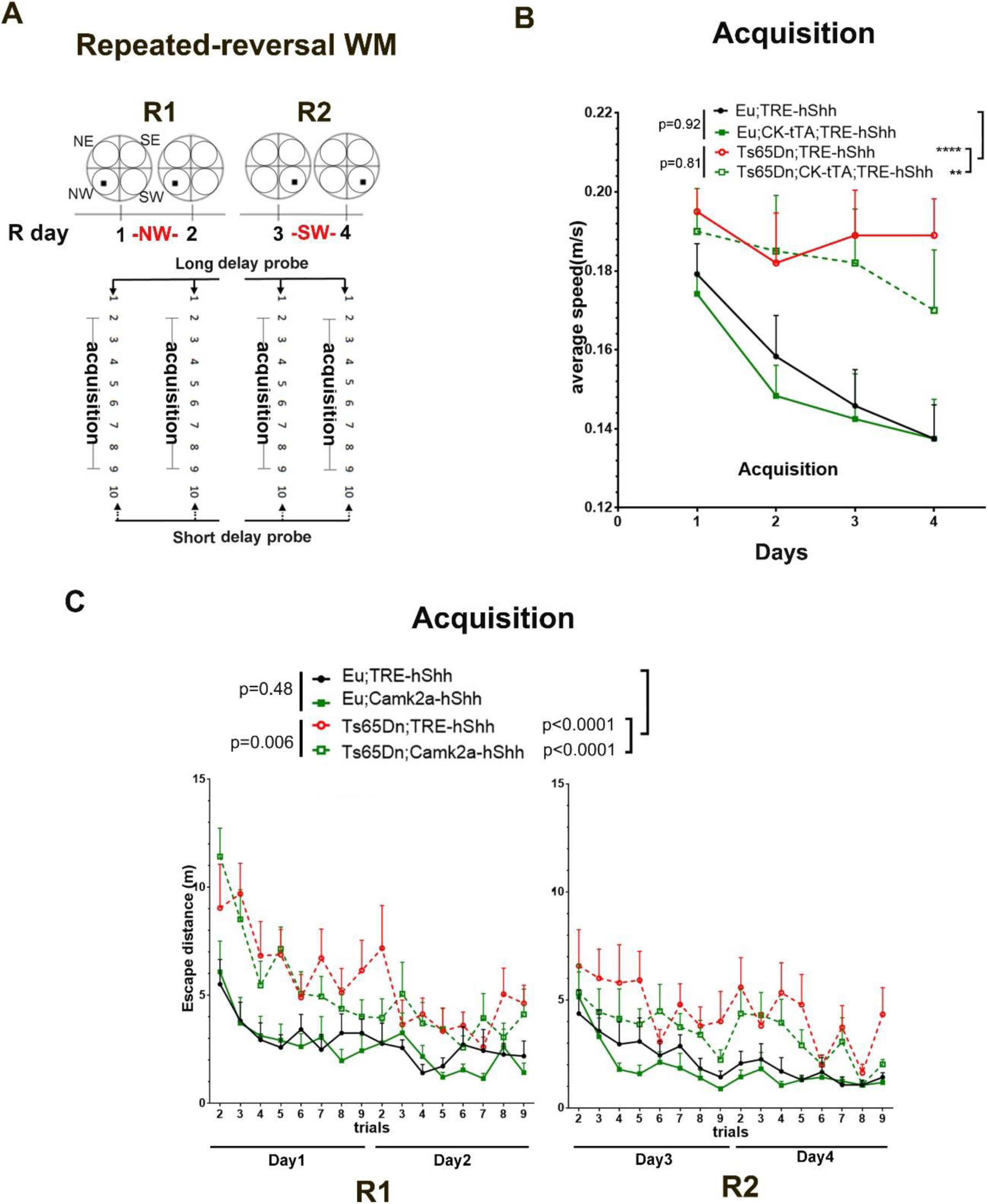
Supplements for RRWM, related to. Figure 6 (A) Scheme illustrating RRWM. (B) Swimming speed in acquisition trials of RRWM. (C) Escape distance in acquisition trials of RRWM. Data are analyzed by two-way ANOVA and Tukey’s multiple comparisons test and expressed as mean ± SEM.

## Supplemental Videos

**Video S1 and S2.** Sagittal brain sections of 2-month-old Camk2a-hShh mice were immunostained with anti-Shh, and the hippocampus region were Z-stack and tile scan imaged with DAPI, Zsgreen, and RFP (anti-Shh) confocal channels, which were converted into movies by Imaris. Two channels, DAPI and Zsgreen, were shown in (Video S1), and all three channels were shown in (Video S2).

**Video S3.** Sagittal brain sections of 2-month-old Pcp2-hShh mice were immunostained with anti-Calbindin1, the cerebellum region were Z-stack and tile scan imaged with DAPI, Zsgreen, and RFP (anti-Calbindin1) confocal channels that were converted into a movie.

## Supplemental Tables

**Table S1. Strategy to test Camk2a-rtTA;TRE-LacZ and Pcp2-rtTA;TRE-LacZ mice**

**Table S2. Experimental animal information**

**Table S3. Oligonucleotides**

**Table S4. Measurements from MRI of the cerebellum cohort**

**Table S5. Consolidated tables for all statistical analysis**

