## Supplementary material for "Forebrain Shh overexpression improves cognitive function in a Down syndrome mouse model and euploid littermates": key sources table

**KEY RESOURCES TABLE**

| **REAGENT or RESOURCE** | **SOURCE** | **IDENTIFIER** |
| --- | --- | --- |
| **Antibodies** | | |
| Shh (C9C5) Rabbit mAb antibody | Cell Signaling Technology | Cat# 2207, RRID:AB_2188191 |
| Sonic Hedgehog antibody [EP1190Y] | Abcam | Cat# ab53281, RRID:AB_882648 |
| SHH Monoclonal Antibody (5H4) | Thermo Fisher Scientific | Cat# MA5-17173, RRID:AB_2538644 |
| GLI1 (L42B10) Mouse mAb antibody | Cell Signaling Technology | Cat# 2643, RRID:AB_2294746 |
| Calbindin (D1\|4Q) XP Rabbit mAB antibody | Cell Signaling Technology | Cat# 13176, RRID:AB_2687400 |
| Anti-β-Actin Antibody | Sigma-Aldrich | Cat# A5441, RRID:AB_476744 |
| Pierce Goat anti-Mouse IgG (H+L) Cross Adsorbed Secondary Antibody, HRP conjugate | ThermoFisher Scientific | Cat# 31432 |
| Pierce Goat anti-Rabbit IgG (H+L) Cross Adsorbed Secondary Antibody, HRP conjugate | ThermoFisher Scientific | Cat# 31462 |
| Goat anti-Mouse IgG (H+L) Cross-Adsorbed Secondary Antibody, Alexa Fluor 568 | ThermoFisher Scientific | Cat# A11004 |
| Goat anti-Rat IgG (H+L) Cross-Adsorbed Secondary Antibody, Alexa Fluor 488 | ThermoFisher Scientific | Cat# A11006 |
| Goat anti-Rabbit IgG (H+L) Cross-Adsorbed Secondary Antibody, Alexa Fluor 488 | ThermoFisher Scientific | Cat# A11008 |
| Goat anti-Rabbit IgG (H+L) Cross-Adsorbed Secondary Antibody, Alexa Fluor 568 | ThermoFisher Scientific | Cat# A11011 |
| Goat anti-Rat IgG (H+L) Cross-Adsorbed Secondary Antibody, Alexa Fluor 568 | ThermoFisher Scientific | Cat# A11077 |
| Goat anti-Mouse IgG1 Cross-Adsorbed Secondary Antibody Alexa Fluor 488 | ThermoFisher Scientific | Cat# A21121 |
| **Bacterial and Virus Strains(N/A)** | | |
| N/A | N/A | N/A |
| **Biological Samples** |  |  |
| N/A | N/A | N/A |
| **Chemicals, Peptides, and Recombinant Proteins** | | |
| Proteinase K | ThermoFisher Scientific | Cat#25530015 |
| Rediprime II DNA Labeling System | Cytiva | Cat#RPN1633 |
| Lipofectamine 3000 Reagent | ThermoFisher Scientific | Cat#L3000008 |
| Doxycycline hyclate | Sigma-Aldrich | Cat#D9891 |
| Doxycycline diets (food pellets, 625mg/kg) | Envigo | TD.01306 |
| Trizol | ThermoFisher Scientific | Cat#15596026 |
| Chloroform | MilliporeSigma | Cat#288306 |
| RNeasy Mini Kit | QIAGEN | Cat#74104 |
| high-capacity cDNA reverse transcription kit | ThermoFisher Scientific | Cat#4368814 |
| TaqMan Gene Expression Master Mix | ThermoFisher Scientific | Cat#4369016 |
| Pierce BCA Protein Assay Kit | ThermoFisher Scientific | Cat#23227 |
| DAPI | MilliporeSigma | Cat#10236276001 |
| ProLong Gold Antifade Mountant with DAPI | ThermoFisher Scientific | Cat#P36931 |
| Xgal | Corning Cellgro | Cat#46-101-RF |
| DPX Mountant for histology slide mounting medium | MilliporeSigma | Cat#06522 |
| Recombinant Mouse Sonic Hedgehog/Shh, N-Terminus Protein | R&D systems | Cat#461-SH-025 |
| Recombinant human Shh-N | PeproTECH | Cat#100-45 |
| Recombinant Human Sonic Hedgehog/Shh Protein(hShh-Np), High Activity | R&D systems | Cat#8908-SH |
| Reporter Lysis 5X Buffer | Promega | Cat#E3971 |
| **Critical Commercial Assays** | | |
| [TARGATT™ Fast & Site-Specific Knockin Mouse Models](https://www.appliedstemcell.com/research/services/targatttm-genome-editing/targatt-fast-site-specific-knock-in-mice-service-c57bl6j-asc-5021t) | Applied StemCell | Cat#ASC-5021T |
| [TARGATT™ Mouse Transgenic Kit](https://www.appliedstemcell.com/research/services/targatttm-genome-editing/targatt-transgenic-kit-ast-1003) | Applied StemCell | Cat#AST-1003 |
| Papain Dissociation System | Worthington-biochem | Cat#LK003153 |
| Beta-Glo Assay System | Promega | Cat#E4720 |
| **Deposited Data** | | |
| Raw and analyzed data | This paper | N/A |
| **Experimental Models: Cell Lines** | | |
| MEF | This paper | N/A |
| **Experimental Models: Organisms/Strains** | | |
| Mouse: B6.Cg-Tg(TetO-hShh,-Zsgreen1), also known as B6.TRE-hShh | This Paper | N/A |
| Mouse: B6C3H-Ts65Dn | The Jackson Laboratory | JAX 001924 |
| Mouse: B6.Cg-Tg(Pcp2-HA/rtTA)1Hirai | Riken | RBRC03711 |
| Mouse: B6.Cg-Tg(Camk2a-rtTA)1237Kndl | EMMA | EM:01151 |
| Mouse: B6.Cg-Tg(Pcp2-tTA)3Horr/J | The Jackson Laboratory | JAX 005625 |
| Mouse: B6.Cg-Tg(Camk2a-tTA)1Mmay/DboJ | The Jackson Laboratory | JAX 007004 |
| Mouse: SJL.Cg-Tg(tetop-lacZ)2Mam/J, also known as TRE-LacZ | Hazzard, Karen (NIH/NHGRI) | PMID 7937760  JAX 002621 |
| Mouse: B6.Cg-Gli1^tm2Alj^/J, also known as Gli1-LacZ | The Jackson Laboratory | JAX 008211 |
| **Oligonucleotides** | | |
| Please see supplemental table S3 | In this Paper | N/A |
| **Recombinant DNA** | | |
| hShh cDNA | DNASU | HsCD00082632 |
| pTRE3G-bi-ZsGreen1 | TaKaRa | Cat#631337 |
| pTRE3G-bi-hShh-Zsgreen1 | This paper | N/A |
| pBS-SK(+) | Agilent | Cat#212205 |
| pBS-SK(+)-TRE-bi-hShh-Zsgreen1 | This paper | N/A |
| pBT378 | Applied StemCell | ASC-5021T |
| pBT378-TRE-bi-hShh-Zsgreen1 | This paper | N/A |
| pROSA26-PA | Addgene | Plasmid #21271 |
| pCMV-Tet3G (pCMV-rtTA) | TaKaRa | Cat#631335 |
| **Software and Algorithms** | | |
| QuantStudio 6 and 7 software | ThermoFisher Scientific | <https://www.thermofisher.com/us/en/home/global/forms/life-science/quantstudio-6-7-flex-software.html> |
| SanpGene | GSL Biotech LLC | <https://www.snapgene.com/> |
| [Fiji - ImageJ](https://imagej.net/Fiji) | N/A | <https://imagej.net/ImageJ> |
| Zen | Zeiss | <https://www.zeiss.com/microscopy/us/products/microscope-software/zen-lite.html> |
| Imaris | Bitplane | <https://imaris.oxinst.com/> |
| FlowJo | BD Biosciences | <https://www.flowjo.com/solutions/flowjo> |
| GraphPad Prism | GraphPad | <https://www.graphpad.com/scientific-software/prism/> |
| Softmouse | Iseehear | <https://www.softmouse.net/> |
| Any-maze | Stoeltingco | <https://www.stoeltingco.com/anymaze.html> |
